# CDK1 phosphorylates ULK1-ATG13 complex to regulate mitotic autophagy and Taxol chemosensitivity

**DOI:** 10.1101/634733

**Authors:** Zhiyuan Li, Xiaofei Tian, Xinmiao Ji, Dongmei Wang, Xin Zhang

## Abstract

ULK1-ATG13 is the most upstream autophagy initiation complex that is phosphorylated by mTORC1 and AMPK to induce autophagy in asynchronous conditions. However, the phospho-regulation and function of ULK1-ATG13 in mitosis and cell cycle remains unknown. Here we show that ULK1-ATG13 complex is differentially regulated throughout the cell cycle. Notably, in mitosis, both ULK1 and ATG13 are highly phosphorylated by CDK1/cyclin B, the key cell cycle machinery. Combining mass spectrometry and site-directed mutagenesis, we found that CDK1-induced ULK1-ATG13 phosphorylation positively regulates mitotic autophagy and Taxol chemosensitivity, and some phosphorylation sites occur in cancer patients. Moreover, double knockout of ULK1 and ATG13 could block cell cycle progression and significantly decrease cancer cell proliferation in cell line and mouse models. Our results not only bridge the mutual regulation between the core machineries of autophagy and mitosis, illustrate the mitotic autophagy regulation mechanism, but also provide ULK1-ATG13 as potential targets for cancer therapy.

## Introduction

Autophagy occurs at basal levels in most tissues to selectively eliminate unwanted cellular components and can also be induced in response to various physiological and pathological conditions. Evolutionarily conserved autophagy-related (ATG) proteins play essential roles in autophagy nucleation, elongation, autophagosome closure and maturation [1–3]. In higher eukaryotes, many ATG proteins have diverse physiologically vital roles not only in autophagy, but also in other *ATG* gene-dependent pathways, such as phagocytosis, secretion and exocytosis, innate immunity and cell cycle etc [2]. Although some ATG proteins such as ATG7, FIP200, Beclin-1 and ATG5 function in cell cycle and mitosis regulation [4–7], whether the most upstream member, serine/threonine UNC-51-like kinase (ULK1/ATG1) [1, 8–11], participates in cell-cycle and mitosis regulation has not been investigated.

On the other hand, although the currently established autophagy regulation mechanisms are mostly from asynchronous cells, in which only around 5% or less are in mitosis, recent studies suggest that autophagy is differentially regulated throughout the cell cycle [12–15], especially in mitosis [16, 17]. Although the autophagosome number at a fixed timepoint is much reduced in mitotic cells compared to interphase cells [16], the autophagic flux is actually active [15, 17, 18]. Moreover, it has been reported that multiple kinases are involved in both autophagy and mitosis [12, 14], indicating that these two cellular processes are intertwined. However, the mitotic autophagy regulation is still under explored.

The only work that has investigated the mitotic autophagy regulation mechanism so far is by Furuya et al [16]. They reported reduced phosphatidylinositol-3-phosphate (PtdIns3P) in mitosis, which suggested decreased VPS34 complex activity. They further identified VPS34-Thr159 to be the mitotic specific- phosphorylation site by CDK1 (the mammalian homolog of Cdc2 in yeast), which is one of the cyclin-dependent kinases (CDKs) that coordinate with their cyclin partners to regulate cell cycle progression. However, whether other molecular mechanisms are involved in mitotic autophagy regulation is still unknown, especially the one that is responsible for the autophagy flux maintenance in mitosis.

Although ULK1-ATG13, the core machinery for the ULK1 autophagy initiation complex, was regulated by phosphorylation primarily from mammalian target-of-rapamycin (mTOR) and adenosine monophosphate activated protein kinase (AMPK) to control autophagy induction in asynchronous cells [8, 9, 19, 20], its regulation mechanism and function in mitosis and the specific cell cycle is uncovered. Here we found that ULK1-ATG13 not only play essential roles in cell cycle progression, but also is directly phosphorylated by CDK1/cyclin B in mitosis to regulate mitotic autophagy and Taxol-induced cell death.

## Results

### ULK1-ATG13 is differentially regulated during cell cycle

To dissect the underlying mechanism of ULK1-ATG13 regulation during cell cycle and mitosis, we synchronized HeLa cells using double-thymidine and nocodazole. Surprisingly, both ULK1 and ATG13 underwent a significant electrophoretic mobility shift in mitosis, while other ATGs such as ATG5, Beclin-1 or ATG101 did not (Figure 1A). In addition, thymidine was used in combination with RO-3306 [21], a specific CDK1 inhibitor, to synchronize cells to specific phases of mitosis. We found that the ULK1 and ATG13 electrophoretic mobility shift were closely correlated with mitotic progression (Figure 1B).

**Figure 1.**
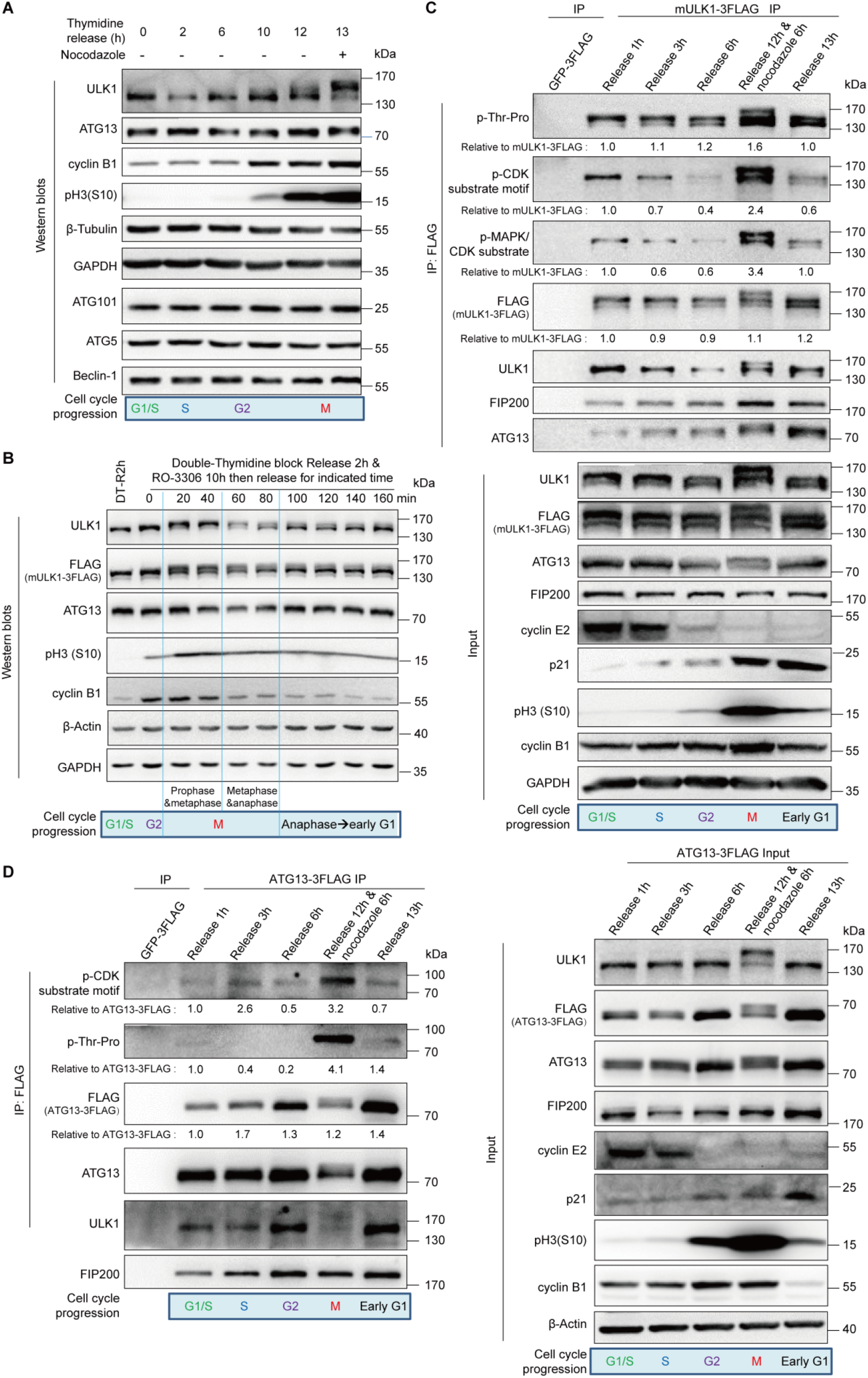
ULK1-ATG13 is differentially regulated during the cell cycle. (A) ULK1-ATG13 shows a mobility shift in mitosis. HeLa cells synchronized by double-thymidine release in the presence or absence of nocodazole were subjected to SDS-PAGE and Western blots analysis. (B) ULK1-ATG13 undergoes band shift during mitotic progression. HeLa cells synchronized by double-thymidine and RO-3306 were released into mitosis for Western blots analysis. (C-D) ULK1-ATG13 upshifted band in mitosis could be recognized by CDK substrate-specific antibodies. 293T cells stably expressing FLAG-tagged mULK1 or ATG13 were synchronized by single-thymidine and released in the presence or absence of nocodazole. The co-immunoprecipitates and input were immunoblotted with specific antibodies. Various cell cycle markers were detected to show the respective phases of cell cycle.

Since phosphorylation-induced mobility shift is often indicative of phosphorylation on serine/threonine-proline residues [22], we seek to perform immunoprecipitation (IP) experiments and examined the ULK1 or ATG13 IP products using the motif antibodies for p-MAPK/CDK Substrate (PXS*P or S*PXR/K), p-CDK Substrate Motif [(K/H)pSP] and phospho-Threonine Proline [23–25]. Since mouse ULK1 has been validated in multiple studies [8, 9, 26–29], We constructed HEK-293T cell lines overexpressing FLAG-tagged mULK1, FLAG-tagged ATG13 or FLAG-tagged GFP control and performed immunoprecipitation in cells after thymidine block release with or without nocodazole (Figures 1C and 1D). Indeed, both the ULK1/FLAG antibodies and the CDK or MAPK/CDK substrate-specific motif antibodies could recognize a significant amount of ULK1 or ATG13 in both upshifted and non-shifted bands in mitotic cells compared with cells in other phases, which further indicates that ULK1-ATG13 was differentially regulated at both protein and phosphorylation levels (Figures 1C and 1D).

### ULK1 is highly phosphorylated in mitosis

To better understand the detailed ULK1-ATG13 regulation mechanisms in mitosis, ULK1 and ATG13 were investigated in detail respectively. The immunoprecipitated ULK1 shows an obvious band shift in mitosis, both on the Coomassie brilliant blue stained gel as well as on Western blots (Figure 2A). Using a phospho-serine/threonine antibody, we confirmed that ULK1 is highly phosphorylated on serine/threonine in mitotic cells compared with asynchronous cells (Figure 2A).

**Figure 2.**
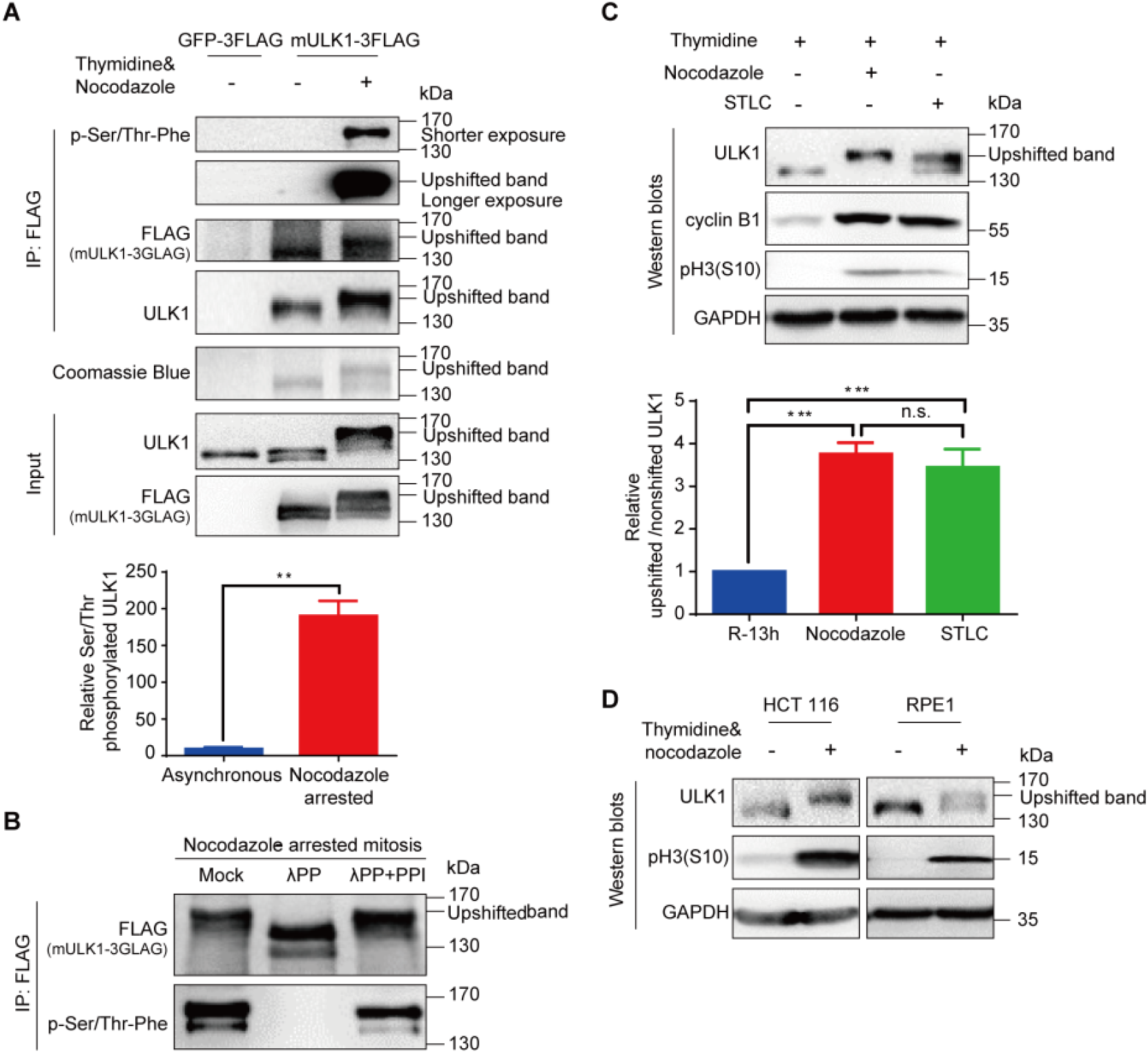
ULK1 is highly phosphorylated in mitosis. (A) ULK1 is phosphorylated in nocodazole-arrested mitosis. 293T cells overexpressing FLAG-tagged mULK1 or GFP were synchronized by single-thymidine and nocodazole. The immunoprecipitates using the FLAG antibody were subjected to Coomassie Brilliant Blue R-250 staining and Western blots analysis. Statistical analysis for relative serine/threonine phosphorylated ULK1 was shown in Figure 2A, lower panel. n=4, **p < 0.01. (B) The immunoprecipitates from Figure 2A were treated with or without λ phosphatase in the presence or absence of phosphatase inhibitors and then subjected to Western blots analysis. λ PP, lambda phosphatase. (C) ULK1 undergoes similar mobility shift in both nocodazole- and STLC-arrested mitosis. HeLa cells synchronized with single-thymidine and nocodazole or STLC were analyzed by Western blots. The upper panel shows the immunoblotting and lower panel shows the ratio of upshifted and non-shifted ULK1. One-way ANOVA followed by Tukey’s Multiple Comparison Test was used for the analysis. n=5, n.s., not significant, ***p <0.0001. (D) ULK1 undergoes phosphorylation-induced electrophoretic mobility shift in single-thymidine and nocodazole synchronized mitotic HCT 116 and RPE1 cells analyzed by Western blots.

To confirm that the band shift of ULK1 in mitosis is due to phosphorylation, we treated the ULK1 immunoprecipitation products with lambda phosphatase in the presence or absence of its inhibitors (Figure 2B). The FLAG-tagged mULK1 band is downshifted by lambda phosphatase treatment, which can be reversed by phosphatase inhibitors. Moreover, the phospho-serine/threonine detected band was also diminished after lambda phosphatase treatment, which can also be reversed by phosphatase inhibitors (Figure 2B). This confirmed that the electrophoretic mobility upshift of ULK1 in mitosis is due to serine/threonine phosphorylation.

To rule out the possibility that the band shift was caused by nocodazole-induced microtubule disruption, but not mitosis *per se*, we used STLC, a specific Eg5 inhibitor [30], to synchronize HeLa cells to mitosis. We found that STLC could induce ULK1 band shift similar to nocodazole treatment, although to a less extent (Figure 2C). The dramatic band shift phenomena were also verified for endogenous ULK1 in HCT 116 (human colorectal cancer cells) and RPE1 (human retinal pigmented epithelial cells) (Figure 2D), as well as endogenous ULK1 and exogenous mouse ULK1 in HEK-293T and HeLa cells with or without FLAG-tagged mULK1 overexpression (Figures S1A and S1B). Therefore, ULK1 is highly phosphorylated in mitosis of multiple cell types.

It should be mentioned that the original antibody we used for ULK1 (Cell signaling technology, #8054) did not detect the upshifted band, but only showed decreased signal in mitosis (Figure S1C, the upper blot). However, it is interesting that when the PVDF membrane was treated with lambda phosphatase to remove the phosphorylation, the upshifted band appeared (Figure S1C, the lower blot), which indicates that the mitotic ULK1 phosphorylation might interfere with the recognition of this specific antibody. Therefore, for all work other than Figure S1C and Figure 3E, we used ULK1 antibody (Cell signaling technology, #4776) instead because it is consistent with the FLAG antibody (detecting the FLAG-tagged mULK1) and Coomassie staining in the FLAG-tagged mULK1 expressing cells (Figure 2A).

**Figure 3.**
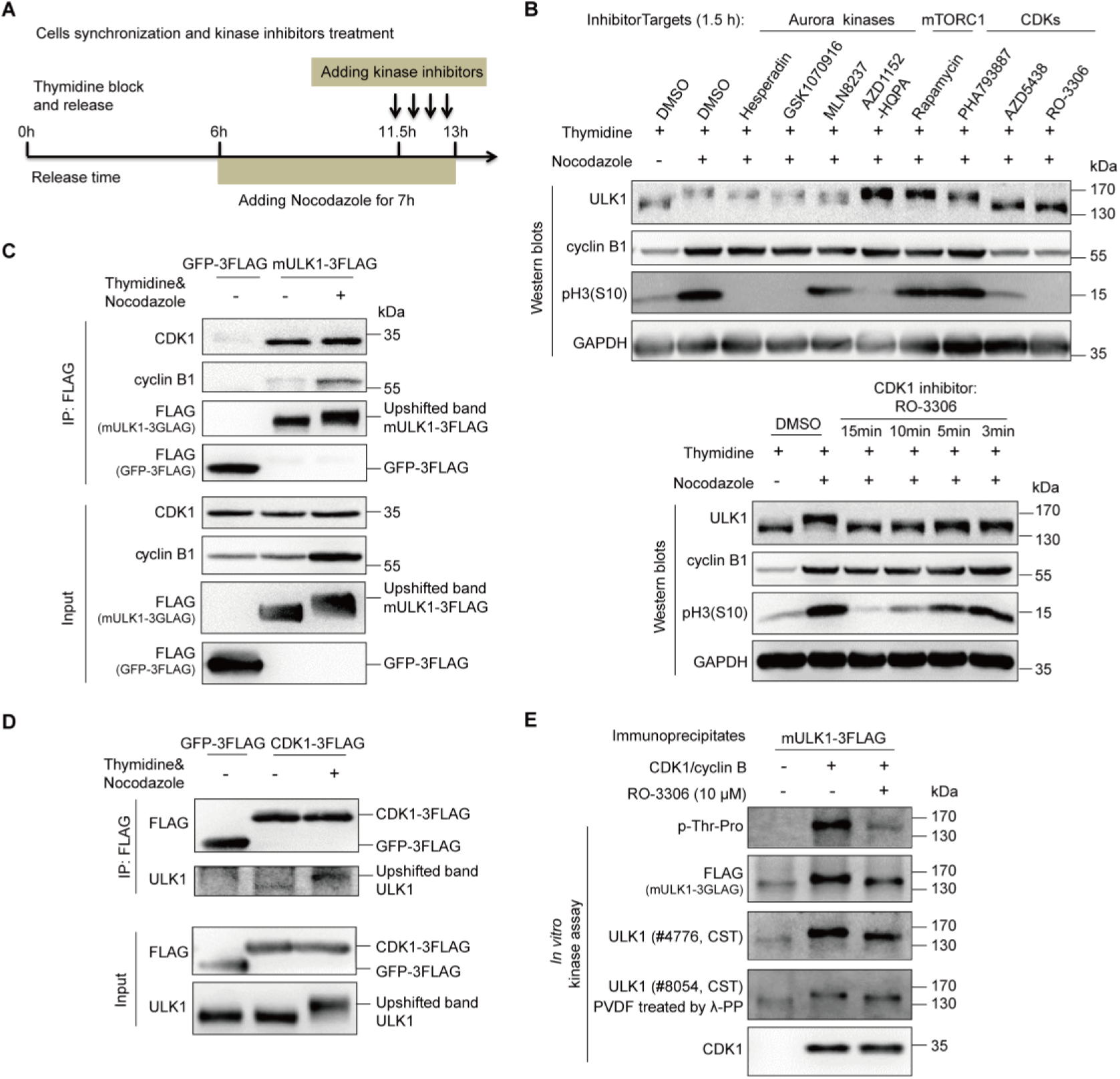
ULK1 is a substrate of CDK1/cyclin B in mitosis. (A) Illustration of cell synchronization and kinase inhibitors treatment. HeLa cells were synchronized with single-thymidine and nocodazole. Kinase inhibitors were added for different timepoints, ranging from 3 min to 1.5 h. (B) CDK1 inhibitors, but not Aurora kinase, mTORC1 and other CDKs inhibitors, abolished the ULK1 bandshift in mitosis. HeLa cells synchronized and treated as (A) were subjected to Western blots analysis. (C-D) ULK1 co-immunoprecipitates with CDK1 and vice versa. 293T cells stably overexpressing FLAG-tagged mULK1 or CDK1 were synchronized with single-thymidine and nocodazole and the co-immunoprecipitates were subjected to Western blots analysis. (E) ULK1 is upshifted and phosphorylated by purified CDK1/cyclin B complex *in vitro*. Purified CDK1/cyclin B complex as kinase and the ULK1 immunoprecipitates from asynchronous 293T cells overexpressing FLAG-tagged mULK1 as substrate with or without RO-3306 were subjected to *in vitro* kinase assay and Western blots analysis. λ-PP, lambda phosphatase.

### ULK1 is a direct substrate of CDK1/cyclin B in mitosis

In an attempt to elucidate the upstream kinase responsible for ULK1 phosphorylation in mitosis, we first used Scansite 3 (http://scansite3.mit.edu/) to predict the potential kinases with proline-directed serine/threonine motif preference, which indicated that CDK1, CDK5, MAPK1/3 are potential candidates. In the meantime, we used a combination of cell synchronization and various kinase inhibitors (Figure 3A), including Aurora kinase inhibitors Hesperadin, GSK1070916, MLN8237 and AZD1152-HQPA [31–34], mTORC1 inhibitor Rapamycin [35] and CDKs inhibitors PHA793887, AZD5438 and RO-3306 [21, 36, 37], to examine their effects on mitotic ULK1 mobility shift (Figure 3B). While none of the inhibitors of Aurora kinases, mTORC1, CDK2, CDK5 or CDK7 could significantly reduce the ULK1 mobility shift, it was interesting that both CDK1 inhibitors AZD5438 and RO-3306 completely abolished the ULK1 mobility shift. In contrast, the CDK2/5/7 inhibitor PHA793887 did not (Figure 3B, the upper panel). This indicates that CDK1 is likely to be the ULK1 upstream kinase responsible for its mobility shift in mitosis.

Considering that 1.5-hour treatment by CDK1 inhibitors could potentially affect the cell cycle themselves, which could in turn affect ULK1 phosphorylation. In fact, we did observe a significant decrease in both pH3(S10) and cyclin B1 levels (Figure 3B, the upper panel), as well as dramatic cell morphological changes with pronounced blebbing, which is consistent with a previous report [21]. Therefore, it is possible that the reduction of mitotic ULK1 mobility shift was simply caused by CDK1 inhibition-induced mitotic exit. Next, to avoid affecting the cell cycle, we treated the mitotic HeLa cells with RO-3306 for as short as 3 minutes, which did not reduce the pH3(S10) or cyclin B1 level (Figure 3B, the lower panel). However, we still observed the significant reduction of mitotic ULK1 mobility shift, which confirmed that CDK1 is indeed the major upstream kinase that phosphorylates ULK1 and induces its mobility shift in mitosis (Figure 3B, the lower panel).

Given that CDK1/cyclin B has multiple substrates [38], we further examined the *in vivo* association between ULK1 and the CDK1/cyclin B kinase complex. ULK1 immunoprecipitation revealed that CDK1 and its mitotic partner cyclin B1 can be co-immunoprecipitated by ULK1 (Figure 3C). In contrast, the other predicted potential kinases by Scansite, such as MAPK1/3 and Aurora A, were not. Reciprocally, we also established a FLAG-tagged CDK1 overexpressing 293T cell line and found that ULK1 could also be co-immunoprecipitated by the FLAG-tagged CDK1 (Figure 3D), which confirmed that CDK1 interacts with ULK1. Importantly, we also performed *in vitro* kinase assay using purified CDK1/cyclin B kinase complex. We found that ULK1 could be highly phosphorylated and upshifted by CDK1/cyclin B complex and the CDK1 inhibitor RO-3306 antagonized its phosphorylation and upshift (Figure 3E), which confirms that ULK1 is a direct substrate of CDK1.

Given that ULK1 could be autophosphorylated due to its kinase activity, the mobility shift and phosphorylation of kinase dead ULK1-K46I mutant [39] in mitosis were investigated. Our results show that ULK1-K46I could undergo mobility shift and mitotic phosphorylation as well (Figures S2A and S2B), which suggest that the kinase dead ULK1-K46I can also be phosphorylated in mitosis. Given that ATG13 is a substrate of ULK1, the ULK1-K46I could not phosphorylate ATG13, which reduced its electrophoresis bandshift and also had a significantly decreased interaction with ATG13 (Figure S2B, the input and IP for ATG13). Consistent with the cellular experiments, *in vitro* kinase assay also suggested that CDK1/cyclin B could phosphorylate ULK1-K46I and induce its upshift (Figure S2C). Additionally, ULK2, a member of ULK1 kinase family, was also found to be phosphorylated in mitosis with CDK1 substrate motif antibody and FLAG antibody (detecting FLAG-tagged mouse ULK2, mULK2-3FLAG) (Figure S3).

### ATG13 is also a direct substrate of CDK1/cyclin B in mitosis

Co-immunoprecipitation was conducted using FLAG-tagged mULK1 and FLAG-tagged GFP as control. As components of the ULK1 complex, ATG13 but not FIP200 showed electrophoretic mobility shift in mitosis (Figure 4A). Similar to ULK1, ATG13 mobility shift in mitosis was also decreased by RO-3306 treatment (Figures 4B and S4A). To further confirm that the ATG13 electrophoretic mobility shift was caused by CDK1 as well, co-immunoprecipitation assay was performed in mitotic cells treated with RO-3306 for different timepoints. Consistent with ULK1, the electrophoresis upshift of ATG13 was also abolished by CDK1/cyclin B specific inhibitor RO-3306 (Figure S4B), which confirms that mitotic ATG13 is also phosphorylated by CDK1.

**Figure 4.**
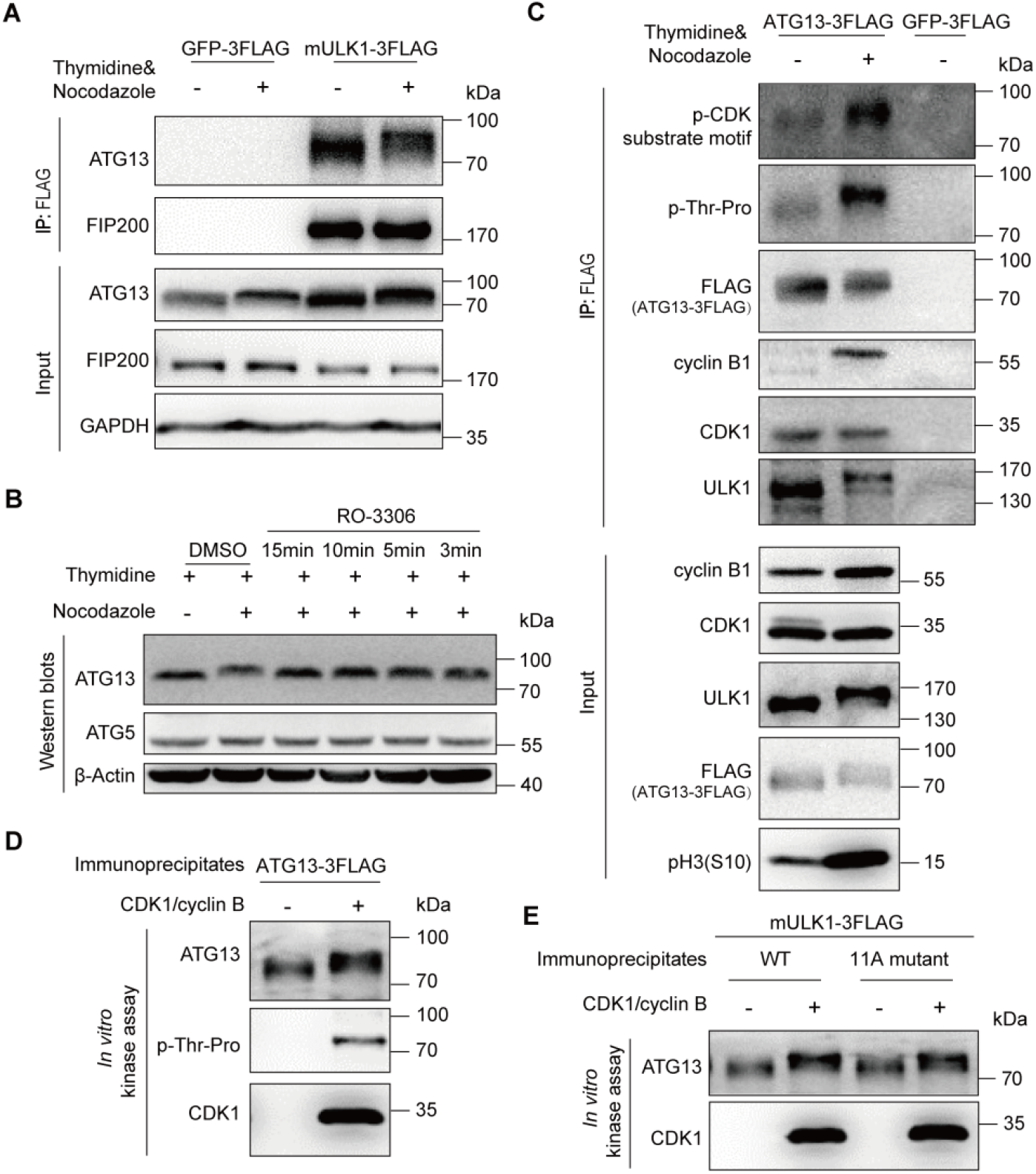
ATG13 is a substrate of CDK1/cyclin B in mitosis. (A) Mitotic ATG13 undergoes mobility upshift in mitosis in electrophoresis. 293T cells overexpressing FLAG-tagged mULK1 or GFP were synchronized by single-thymidine in the presence or absence of nocodazole and co-immunoprecipitated by the FLAG antibody followed by Western Blots. Size of endogenous ATG and expressed FLAG-tagged ATG13 were not distinguishable in electrophoresis here. (B) CDK1 inhibitor RO-3306 decreases the ATG13 bandshift in mitosis. HeLa cells synchronized and treated as Figure 3A were subjected to Western blots analysis. (C) ATG13 is phosphorylated and interacts with CDK1/cyclin B1 in mitosis. 293T cells overexpressing FLAG-tagged ATG13 were synchronized by single-thymidine in the presence or absence of nocodazole. The co-immunoprecipitates by FLAG antibody were subjected to Western blots analysis. (D-E) ATG13 is directly phosphorylated by purified CDK1/cyclin B complex *in vitro*. The ATG13 immunoprecipitates from asynchronous 293T cells overexpressing FLAG-tagged ATG13 or ULK1 and purified CDK1/cyclin B complex were subjected to *in vitro* kinase assay followed by Western blots analysis. (D) shows representative Western blots of the ATG13 immunoprecipitates as substrate, and (E) shows the representative Western blots of the ULK1-WT/11A-mutant (identified in Figure 5E and 5F) co-immunoprecipitates as substrate.

Next, we performed co-immunoprecipitation to examine the interaction between ATG13 and ULK1, CDK1 in 293T cells overexpressing FLAG-tagged ATG13. Both Proline directed phospho-serine and threonine motif antibodies showed significant signal increase in mitosis (Figure 4C). The electrophoretic mobility shift of FLAG-tagged ATG13 was also apparent in mitosis. Moreover, both CDK1 and cyclin B1 could be co-immunoprecipitated by ATG13 (Figure 4C). These results confirmed the interactions among ATG13, ULK1 and CDK1/cyclin B1.

It should be mentioned that in mitosis the ATG13 band shift are indistinguishable between ULK1-knockout and ULK1-overexpression cell lines, but are abrogated by RO-3306. This indicates that ATG13 phosphorylation-associated electrophoretic mobility shift is dependent on CDK1/cyclin B kinase activity but not its well-known kinase ULK1 (Figure S4C). Noteworthy, given that ATG13 could be phosphorylated by ULK1, despite in asynchronous condition, cells with ULK1 knockout showed downshifted band for ATG13. However, ATG13 band shift did not show any change in asynchronous cells with or without ULK1 after RO-3306 treatment (Figure S4D). In fact, *in vitro* kinase assay demonstrates that ATG13 could be directly phosphorylated and upshifted by purified CDK1/cyclin B kinase complex, which are shown by upshifted/phosphorylated ATG13 in both ATG13 immunoprecipitates and ULK1 co-immunoprecipitates (Figures 4D and 4E).

### ULK1 and ATG13 phosphorylation sites in mitosis by CDK1/cyclin B

To identify the ULK1 phosphorylation sites in mitosis, we combined Scansite prediction and mass spectrometry analysis. We first selected three potential sites of mouse derived ULK1, serine 622, threonine 635 and threonine 653, because they had high scores in Scansite prediction for Proline-dependent serine/threonine kinase group (Pro_ST_kin), or their phosphorylation signals were specifically identified in mitotic cells by mass spectrometry (Figure 5A). Site-directed mutagenesis was used to mutate these sites from serine/threonine to unphosphorylatable alanine in FLAG-tagged mULK1. Using 293T cells stably expressing FLAG-tagged GFP as control, immunoprecipitation for FLAG-tagged mutant mULK1-S622A/T635A/T653A was conducted in both asynchronous and mitotic cells. Although none of the single mutations obviously disrupted ULK1 phosphorylation or band shift in mitosis (Figure 5B), S622A&T635A&T653A triple mutant significantly decreased the electrophoretic mobility shift and the phospho-threonine-proline signals compared with wild-type ULK1 (Figure 5C).

**Figure 5.**
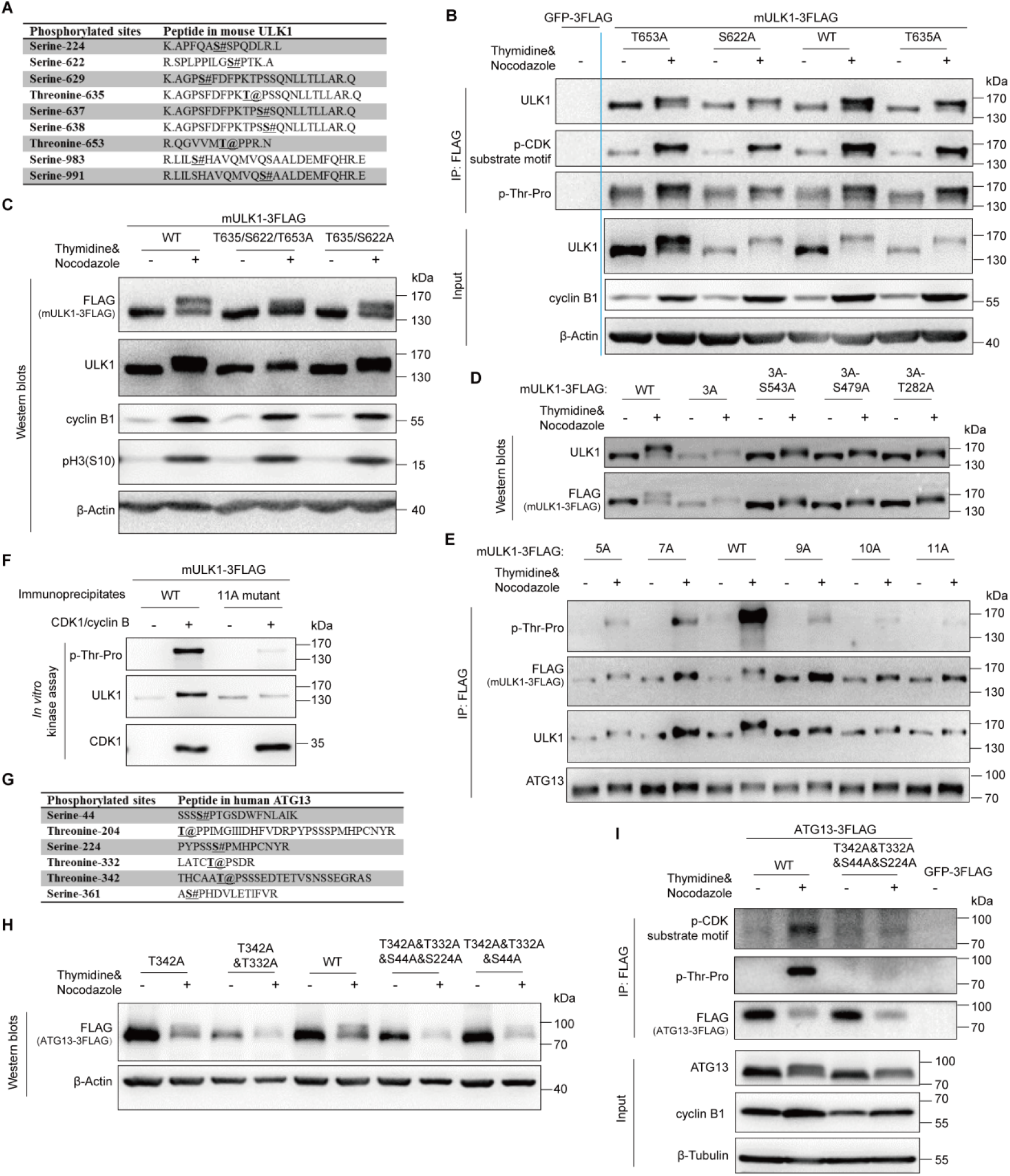
ULK1 and ATG13 phosphorylation sites in mitosis by CDK1/cyclin B. (A) The specific phosphorylated sites identified by mass spectrometry in mitotic mULK1 compared with asynchronous mULK1. (B-C) S622/T635/T653 phosphorylation contributes to ULK1 mobility shift in mitosis. 293T cells overexpressing FLAG-tagged mULK1-S622/T635/T653 mutants were synchronized with single-thymidine and nocodazole. Then immunoprecipitation for single mutant with FLAG antibody (B) or Western blots for double and triple mutant (C) was performed. (D-E) More sites contribute to ULK1 band shift. Based on triple S622/T635/T653A mutant (3A), the other 8 Ser/Thr sites were mutated into Ala. 293T cells expressing various mutants were synchronized into mitosis with thymidine and nocodazole for immunoprecipitation by FLAG antibody and Western blots. 5A, 3A-S479&S543A; 7A, 5A-S411&S413A; 9A, 5A-S413&T401&S403&S405A; 10A, 9A-T282A; 11A, 10A-T502A. (F) ULK1-11A mutant was not upshifted or phosphorylated by CDK1/cyclin B kinase complex *in vitro*. The ULK1-11A mutant immunoprecipitates from asynchronous 293T cells overexpressing FLAG-tagged mULK1 and purified CDK1/cyclin B complex were tested in an *in vitro* kinase assay and Western blots. (G) The phosphorylated sites shared by mass spectrometry identification and Scansite prediction in mitotic ATG13 compared with asynchronous ATG13. (H-I) ATG13-T342/T332/S44/S224 phosphorylation contributes to ATG13 mobility shift in mitosis. 293T cells overexpressing FLAG-tagged ATG13-T342/T332/S44/S224 mutants were synchronized with single-thymidine and nocodazole. Then Western blots for four-site mutant (H) or immunoprecipitation for mutants with FLAG antibody (I) was performed.

Since even the triple mutant did not completely abolish the band shift, we further mutated more sites in the triple mutant (3A) background. It shows that the 11A (S622&T635&T653&S479&S543&S413&T401&S403&S405&T282&T502A) mutant collapsed the electrophoretic mobility shift and abolished ULK1 phosphorylation (as indicated by the p-Thr-Pro signal in Figure 5E) in mitosis, while the other mutants, including the 5A, 7A, 9A and 10A, still have detectable phosphorylation signals and upshift in mitosis (Figures 5D and 5E). It should be mentioned that, as a substrate of ULK1, both ATG13 mobility shift and its interaction with the ULK1-mutant were not much affected by ULK1 mutations, which indicates that the kinase activity of ULK1 is not affected by 11A mutations (ATG13 Western blot in Figure 5E) when compared to the decreased ATG13 interaction with the kinase dead ULK1-K46I (Figure S2B). Importantly, *in vitro* kinase assay using purified CDK1/cyclin B proteins also showed that the ULK1-11A mutant almost completely abolished the phosphorylation signal and band shift compared with wild-type ULK1 (Figure 5F). Therefore, besides the three major sites (S622A/T635A/T653A), CDK1/cyclin B phosphorylates multiple other sites of ULK1 in mitosis.

As for ATG13, we also combined Scansite prediction and mass spectrometry analysis to identify its potential phosphorylation sites in mitosis. We selected four sites, ATG13-T342/T332/S44/S224, that were identified by both Scansite prediction and mass spectrometry analysis (Figure 5G). Site-directed mutagenesis, Western blots and immunoprecipitation were combined for site verification (Figures 5H and 5I), and we found that the four-site mutant significantly decreased the electrophoretic mobility shift and the phospho-serine/threonine-proline signals compared with wild-type ATG13 (Figure 5I).

### ULK1-ATG13 phosphorylation in mitosis positively regulates mitotic autophagy and Taxol-induced cell death

To study the functional significance of these phosphorylation sites, we constructed ULK1 and ATG13 wild-type or double mutant cell line based on HeLa-ULK1&ATG13-DKO cell line (Figures S5A and S5B). Although ULK1-ATG13 complex is a key autophagy regulator, its function has only been investigated in asynchronous cells. To unravel the function of these phosphorylation events that occur specifically in mitosis, we first examined the autophagic flux in mitotic HeLa-DKO cells expressing ULK1&ATG13 WT or mutant cell line. We used autophagic flux, one of the most reliable methods to examine the amount of autophagic degradation to indicate autophagy activity. Autophagic flux inhibitors such as chloroquine (CQ) could cause autophagy marker LC3-II accumulation due to blocked LC3-II degradation [3]. In fact, we found that the autophagic flux was reduced by about 30% in mutant cell lines compared to the wild-type cell lines (Figure 6A), which indicates that ULK1-ATG13 phosphorylation positively regulate autophagy in mitosis.

**Figure 6.**
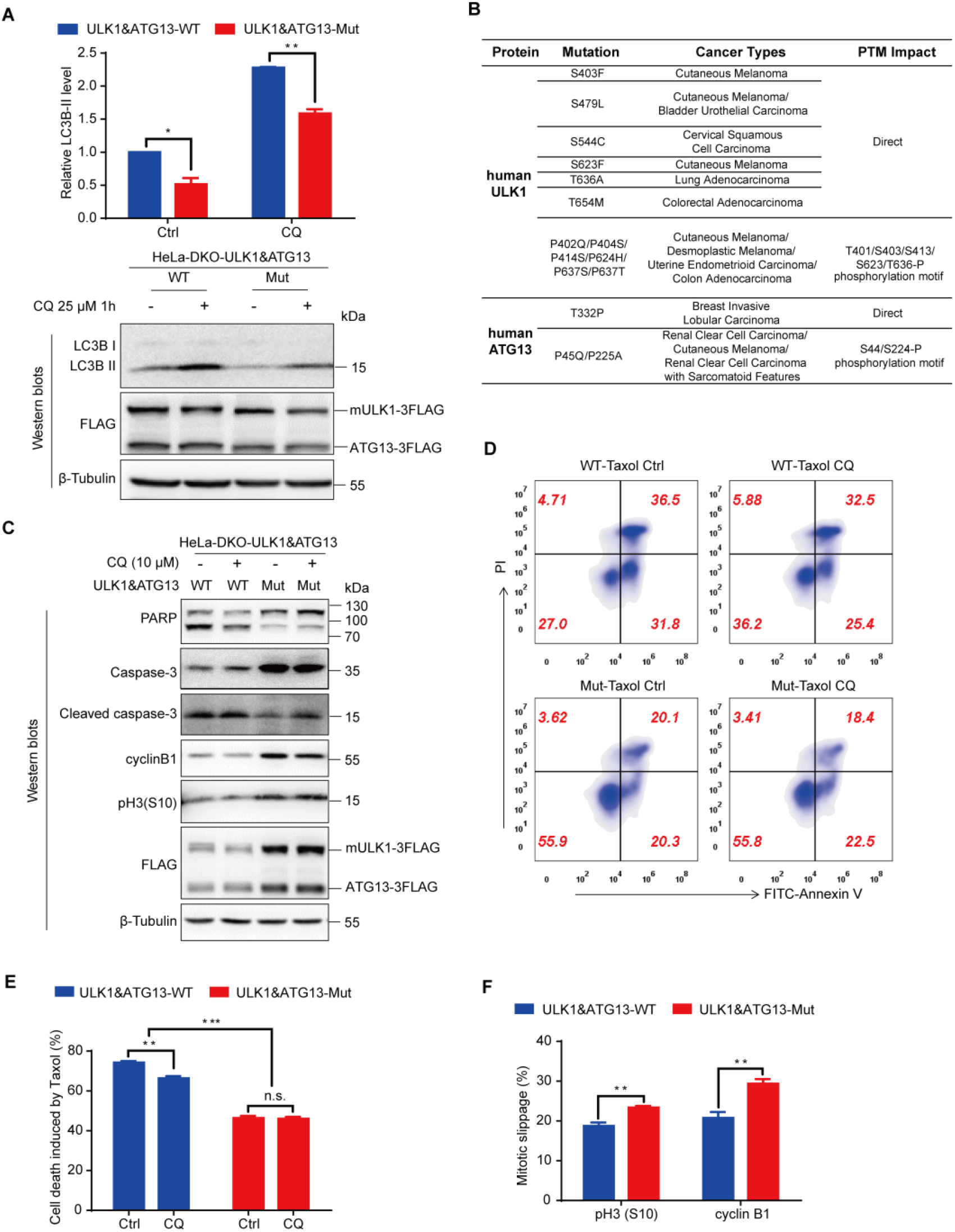
ULK1-ATG13 phosphorylation in mitosis regulates mitotic autophagy and Taxol chemosensitivity. (A) The autophagic flux of HeLa-DKO cells stably overexpressing double wild-type or mutant FLAG-tagged mULK1 and ATG13. Cells synchronized to mitosis with thymidine and nocodazole were shaken off and treated with or without autophagy inhibitor 25 μM CQ for 1h. Western blots and statistical analysis for autophagy marker LC3B-II. The upper panel is the statistical result and the lower panel is a representative Western blot. n=3, *p < 0.05, **p < 0.01. (B) Human ULK1 and ATG13 mutations in cancer patients. The data were extracted from the cBioPortal for Cancer Genomics and arranged according to phosphorylation sites identified in mitotic mouse ULK1 and human ATG13 (Figure 5). It should be pointed out that the human ULK1 mutation sites of S544, S623, T636 and T654 in Figure 6B are equivalent to mouse ULK1 sites of S543, S622, T635 and T653. PTM, Post-translational modification. (C) Analysis of cells treated with Taxol combined with CQ by Western blots. HeLa-DKO cells stably overexpressing double wild-type or mutant FLAG-tagged mULK1 and ATG13 cells released for 4h from thymidine block were treated with 20 nM Taxol with or without 10 μM CQ for 24 h and analyzed by Western blots for apoptosis and mitotic/slippage markers. (D) The representative flow cytometry result in cells as treated in Figure 6C. Cells treated as in Figure 6C were subjected to co-staining with FITC-Annexin V and PI and analyzed by flow cytometry. (E) The statistical analysis for cell death induced by Taxol and/or CQ for Figure 6D. n=3, n.s., not significant, **p < 0.01, ***p < 0.001. (F) Statistical analysis for mitotic slippage markers cyclin B1 and pH3(S10) in cells as treated in Figure 6C. Cells fixed with −20°C 75% Ethanol overnight were subjected to either PI and pH3(S10) co-staining or cyclin B1 staining for cell cycle, mitotic index and cyclin B1 level analysis by flow cytometry. n=3, **p < 0.01.

It is very interesting that some ULK1/ATG13 mutations in patients were found to be associated with the phosphorylation motif/sites identified in our study. For example, in Desmoplastic Melanoma, Cutaneous Melanoma, Uterine Endometrioid Carcinoma and Colon Adenocarcinoma patients, there are some proline mutations such as P402Q/P404S/P414S/P624H/ P637S/P637T in the CDK1 substrate motifs in ULK1 [40, 41], which is likely to indirectly affect CDK1 phosphorylation on ULK1. There are also some direct serine/threonine mutations such as S403F, S479L, S544C, S623F, T636A and T654M in Cutaneous Melanoma, Bladder Urothelial Carcinoma, Cervical Squamous Cell Carcinoma, Lung Adenocarcinoma and Colorectal Adenocarcinoma patients [41–46], which could directly affect CDK1 phosphorylation on ULK1. Similarly, in Renal Clear Cell Carcinoma, Cutaneous Melanoma and Renal Clear Cell Carcinoma with Sarcomatoid Features patients, there are also mutations of P45Q and P225A in the CDK1 substrate motifs in ATG13[41, 47, 48], which is likely to indirectly affect CDK1 phosphorylation on ATG13. And the mutation for ATG13-T332P in Breast Invasive Lobular Carcinoma [49] could directly affect CDK1 phosphorylation on ATG13 (Figure 6B).

Besides the basic scientific question about autophagy regulation in mitosis, more importantly, it has been shown that the combinatorial therapy by autophagy inhibitors and chemotherapeutic drugs arresting cells in mitosis could have better anti-tumor efficacy [50–52]. Next we combined CQ and chemotherapeutic drug Taxol, a reagent stabilizing the microtubule polymer to arrest cells in mitosis [53]. Although neither ULK1-11A nor ATG13-4A alone has significant effects on Taxol chemosensitivity (Figures S6A and S6B), Taxol-induced cell death was substantially attenuated in ULK1&ATG13 double mutant cell line compared with wild-type cell line (Figures 6C-6E). This decreased chemosensitivity in double mutant cell line may be due to its low autophagy activity, which is consistent with previous report that autophagy inhibition could inhibit cell death in mitotic arrest [18]. In addition, autophagy inhibition by CQ could attenuate Taxol chemosensitivity in wild-type cell line (with higher autophagy), but could not further decrease that of the double mutant cell line (with lower autophagy), which indicates that ULK1-ATG13 is the major autophagy pathway for Taxol-induced cell death in mitosis (Figures 6C-6E). Moreover, the double mutant cell line displayed lower mitotic slippage induced by Taxol (Figure 6F), which may be an alternative mechanism for their differential chemosensitivity. Considering the lower Taxol chemosensitivity in ULK1&ATG13 double mutant cell line, our results indicate that patients with similar mutations in both ULK1 and ATG13 might have increased chemotherapy resistance to Taxol.

## ULK1-ATG13 is required for cell cycle progression

To examine the cell cycle regulation by ULK1-ATG13, we constructed ULK1-knockout (KO) HeLa (human cervical cancer cells) and 293T (human embryonic kidney cells) cells using CRISPR/Cas9 (Figures S7A and S7B). ULK1-knockout cells had similar cell cycle distribution compared to wild-type (WT) cells (Figure S7C). To further examine the effect of ULK1 on cell cycle progression, we synchronized the cells by double-thymidine and found that the S/G2 transition in ULK1-knockout cells was slightly delayed compared to wild-type cells (Figure S7D). Given that the G2 and M phases are not distinguishable by propidium iodide (PI) staining alone, we further used pH3(S10), one of the most commonly used mitotic markers [54, 55], to examine whether ULK1 functions in G2/M transition. Although no differences were detected for the G2/M percentage in wild-type and ULK1-knockout cells, the mitotic progression is significantly decreased in ULK1-knockout cells synchronized with thymidine and nocodazole, a microtubule destabilizing reagent. It was shown by the percentage of pH3(S10) positive cells using flow cytometry (Figure S8A) or Western blots analysis for cell cycle markers (Figure S8B). The antibody for p-CDK Substrate Motif [23] recognizes the substrate of CDK, whose phosphorylation level reflects the CDK activity and is used as a mitotic marker. And either the upshifted band of Myt1 or the lower phosphorylation level of Cdc2-Y15 in mitosis [56] could also be used as mitotic progression markers. Given that ULK1 is a serine/threonine protein kinase, the contribution of ULK1 kinase activity to mitotic progression is examined in cell lines expressing wild-type ULK1 or kinase dead ULK1-K46I mutant [8], which indicates that the ULK1 kinase activity has little effect on mitotic progression (Figure S8C).

As the ULK1 partner in ULK1-ATG13 complex, ATG13 was reported to function in mitotic catastrophe [57], ATG13-knockout cells were established using CRISPR/Cas9 as well (Figure S9A). Cell cycle analysis indicated that ATG13 knockout inhibited G2/M transition and decreased the mitotic index (Figures S9B and S9C), indicating that ATG13 is involved in cell cycle regulation. Noteworthy, it has been reported that ATG13 is required for the kinase activity of ULK1 [26], the ULK1 kinase activity is not necessary for mitotic progression (Figure S8C). Therefore, the phenotype of the ATG13-knockout cells is due to the lack of ATG13 rather than the impairment in the kinase activity of ULK1.

To further investigate the role of ATG13 and ULK1 in cell cycle regulation, we combined ULK1 knockout with ATG13 gRNA transient transfection and found that the cell cycle progression was inhibited (Figures S9D and S9E). Then we constructed a ULK1 and ATG13 double knockout (DKO) cell line (Figure S9F) and found that both S/G2 and G2/M transitions were severely delayed (Figures 7A and S9G), while ULK1 or ATG13 knockout alone (Figures S8A and S9B) did not simultaneously delay both S/G2 and G2/M transitions. The immunoblotting of cell cycle markers verified the mitotic index and cell cycle distribution data (Figure 7B). To rule out the off-target effect, rescue assays were performed in knockout cells with exogenously expressed ULK1-ATG13, which confirmed the specificity of ULK1 and ATG13 knockout (Figure S10). Furthermore, the effect of ULK1 and/or ATG13 on mitotic exit markers was analyzed, which shows that DKO interfered with cyclin B1 and pH3 (S10) phosphorylation decrease more significantly than ULK1 or ATG13 single knockout (Figure 7C). Accordingly, the growth rate of HeLa-DKO cells was significantly slowed down compared with HeLa cells (Figure 7D).

**Figure 7.**
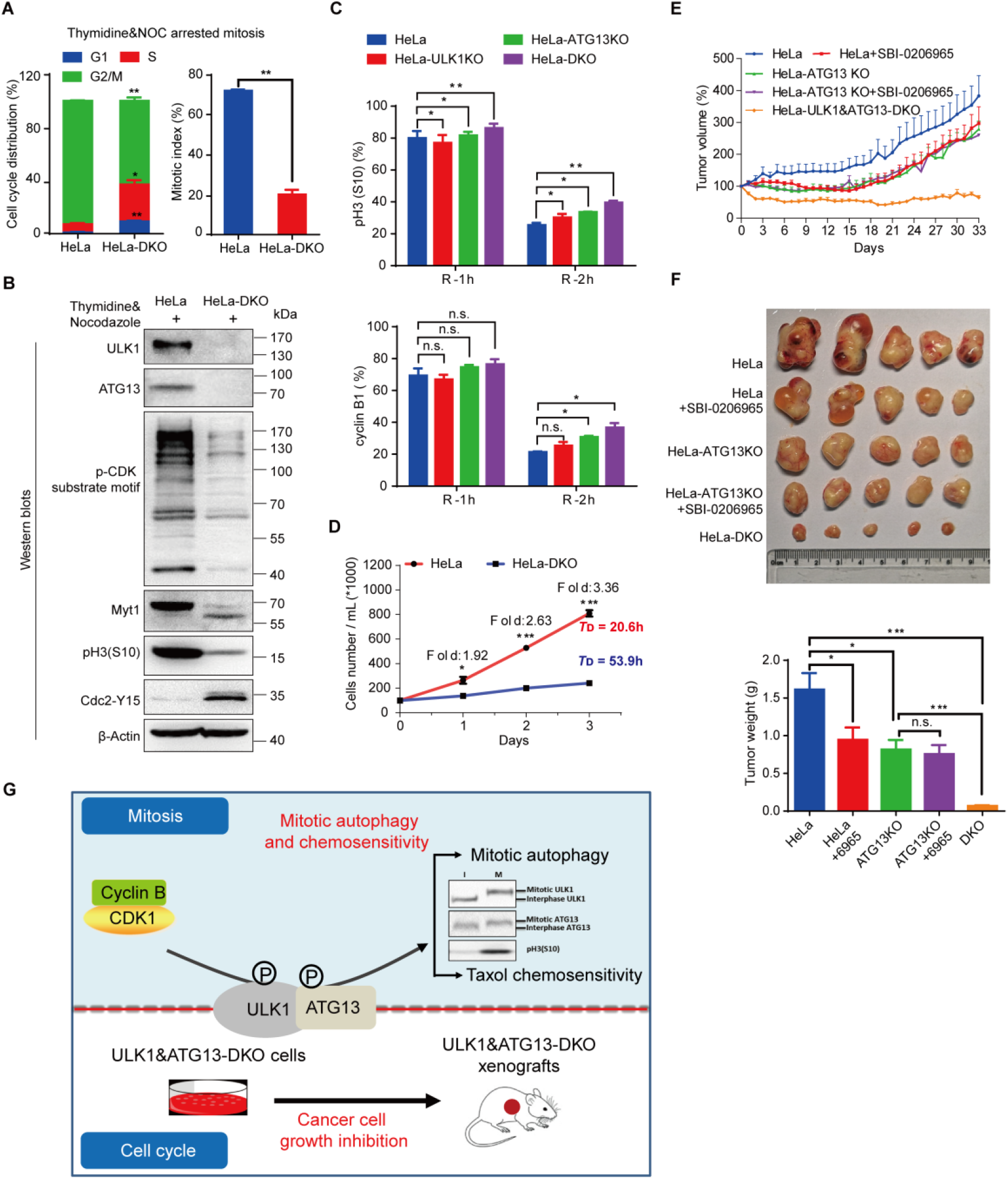
ULK1-ATG13 is required for cell cycle progression. (A) ULK1-ATG13 double knockout (DKO) inhibits S/G2 and G2/M transitions. HeLa wild-type or ULK1&ATG13-DKO cells synchronized into mitosis were subjected to PI and pH3(S10) co-staining for cell cycle and mitotic index analysis by flow cytometry. n=3, *p < 0.05, **p < 0.01. (B) Representative Western blots suggest that ULK1&ATG13 DKO inhibits mitotic entry, which is shown by mitotic markers and CDK1 substrate phosphorylation. (C) Mitotic exit of ULK1, ATG13, or ULK1&ATG13 knockout cells. Cells were synchronized into mitosis with thymidine and nocodazole and released into nocodazole-free complete DMEM medium for different timepoints and then subjected to either PI and pH3(S10) co-staining or cyclin B1 staining for cell cycle, mitotic index and cyclin B1 level analysis by flow cytometry. n=3, n.s., not significant, *p < 0.05, **p < 0.01. (D) ULK1&ATG13 DKO inhibits cell proliferation. HeLa wild-type or ULK1&ATG13 DKO cells were plated at 1×10^5^ cells/ml and cultured for 1, 2 or 3 days. The cells number was counted by flow cytometry and the doubling time was calculated. *T*_D_ indicates the average cell doubling time and is calculated as: *T*_D_= t_*_[lg2/(lgNt-lgN0)], where t is the culture time, Nt is the cell number after culturing, N0 is the original cell number plated. n=3, *p < 0.05, ***p < 0.001. (E) The relative tumor volume growth curve of nude mice bearing different tumors with or without SBI-0206965. The nude mice were injected with cells (1 × 10^7^) in 100 μL PBS/Matrigel Matrix (1:1). Seven days post implantation, five mice in each group were injected with 0.5% (M/V) methyl cellulose or SBI-0206965 in 0.5% methyl cellulose (20 mg/kg/d) every day for 33 days. Tumor growth was evaluated every day and tumor volume was calculated as: volume = 1/2(length × width^2^). (F) Tumor growth in nude mice bearing wild-type or knockout cells with or without ULK1 kinase inhibitor SBI-0206965. The protocols were indicated as Figure 7E, and the mice were sacrificed and tumors were harvested and weighted up at the end of the experiment. n=5, n.s., not significant, *p < 0.05, ***p < 0.001. (G) Model illustrates functions of ULK1-ATG13 in cell cycle and its phospho-regulation in mitosis. When cells are in mitosis, CDK1/cyclin B phosphorylates ULK1-ATG13 to induce significant electrophoretic mobility shift. The phosphorylated ULK1-ATG13 regulates mitotic autophagy and Taxol chemosensitivity. In asynchronous conditions, double knockout ULK1 and ATG13 inhibits cancer cell proliferation in both cell and mouse models. I, interphase; M, mitosis.

To further test the effect of ULK1-ATG13 *in vivo*, the mouse models in nude mice bearing ULK1 and ATG13 single or double knockout cells were established (Figures S11A and S11B). The tumor weight and volume of DKO group were significantly lower than all the other groups (Figures 7E and 7F, Figure S11A). In contrast, ULK1 inhibitor SBI-0206965 [10] and/or ATG13-KO alone were not strong enough to inhibit tumor growth as efficiently as DKO, which proved the synergistic effects for ULK1 and ATG13 (Figures 7E and 7F), indicating that targeting ULK1 and ATG13 might be a potential anti-cancer strategy.

## Discussion

ULK1-ATG13 complex is mainly phosphorylated by AMPK and mTORC1 in asynchronous conditions [8, 9, 19, 20], but little is known about its regulation in mitosis. We found that the master cell cycle kinase CDK1 phosphorylates ULK1-ATG13 complex to regulate its function in autophagy and Taxol chemosensitivity. Besides the known function in autophagy, we also found that ULK1 and ATG13 coordinate to orchestrate cell cycle progression in cell line and mouse models (Figure 7G).

### Kinases involved in both autophagy and mitosis

Recent literature indicates that there are some kinases that are involved in both autophagy and mitosis, which could bridge the autophagy and cell cycle regulation. Cell cycle kinases such as CDK1, Aurora A, PLK-1 were found to regulate autophagy, while kinases originally found in autophagy control were shown to regulate mitosis as well, such as mTORC1 and AMPK [12, 14]. In addition, increased evidence shows that some other autophagic proteins such as Beclin-1, SQSTM1/p62 and GABARAP are related to mitotic events [6, 58–60]. Although it has been reported that ULK3, a member of ULK1 kinase family, could regulate cytokinesis and ATG13 could regulate Colchicinamide-induced mitotic catastrophe [57, 61], the roles of autophagy kinase complex ULK1-ATG13 in mitotic regulation are still unclear. While previous reports implicated possible links between ULK1 and CDK1 [62, 63], our study here is the first report that demonstrates CDK1 is the upstream kinase for ULK1-ATG13 complex. ULK1-ATG13 phosphorylation could positively regulate autophagy, which promoted Taxol-induced cell death and mitotic slippage. In our opinion, the mitotic slippage may be an alternative mechanism for mitotic ULK1-ATG13 phosphorylation mediated autophagy and chemosensitivity (Figure 6). As to the role of ULK1-ATG13 in tumor proliferation, we propose that it is the function of ULK1-ATG13 protein themselves rather than mitotic phosphorylation (Figures 7E and 7F).

### CDK1 interacts with ULK1/ATG13 in asynchronous conditions

It is interesting that CDK1 could be detected in ULK1/ATG13 Co-IP products in asynchronous cells just as in mitotic cells, which indicates the constitutive interaction between CDK1 and ULK1/ATG13. Since Cdc37 is a molecular chaperone linking ULK1-ATG13 to Hsp90, similarly, it is possible that CDK1 may also function as a molecular chaperone linking ULK1-ATG13 complex to Hsp90 or other unknown proteins in interphase [64, 65], which may be important for ULK1-ATG13 stability. CDK1 regulates the S/G2 and G2/M transitions via sequential coupling to cyclin A and cyclin B [66], indicating that ULK1-ATG13 complex could be regulated by CDK1 in both S/G2 and G2/M transitions, which is consistent with the role of ULK1-ATG13 complex and their differential regulation in cell cycle progression (Figures 1 and 7). Obviously, the functional significance for the CDK1 and ULK1/ATG13 interaction in asynchronous conditions, whether other components of ULK1 complex or other ATG proteins are subjected to similar regulation, need further investigations, which would provide more insights for the molecular links between autophagy and cell cycle progression.

### VPS34- and ULK1-ATG13 complex-dependent mitotic autophagy regulation

Although increasing evidence indicates that autophagy in mitosis remains active in mitosis but the regulation mechanism still unclear. Yuan’s group indicated that activated CDK5 phosphorylates VPS34-Thr159 to inhibit VPS34-dependent autophagy in mitosis [16]. Our finding here demonstrates that CDK1 phosphorylates ULK1-ATG13 in mitosis to promote ULK1-ATG13-dependent autophagy, which at least partially contributed to the active autophagy state in mitosis as reported in our previous study [15]. However, it is likely that multiple autophagy regulators, not limited to the VPS34 complex and ULK1-ATG13 complex, contribute to the mitotic autophagy regulation, which certainly needs further investigations. In fact, we also found that ULK2, the homolog of ULK1, is also phosphorylated in mitosis as ULK1 (Figure S3). And ULK1/ULK2-ATG13 complex in mitosis might orchestrate respective functions for autophagy, chemosensitivity and beyond.

### Mitotic ULK1/ATG13 and beyond

The upshifted mitotic ULK1/ATG13 could be viewed as a hint for the other autophagy related proteins potentially functioning in mitosis or cell cycle. Furthermore, deciphering the mitotic phosphatase responsible for ULK1-ATG13 dephosphorylation should also be investigated in the future, which will be complementary for CDK1-mediated phosphorylation. Importantly, controlling the phosphorylation for ULK1-ATG13 by the candidate phosphatases and kinases might be a promising therapeutic strategy for autophagy and mitosis related diseases.

In conclusion, we have uncovered the untraditional roles of ULK1-ATG13 complex in cell cycle progression and tumor growth, which provides ULK1-ATG13 as potential candidates in cancer therapy. We have also revealed its phospho-regulation by CDK1/cyclin B in mitosis, which provides molecular mechanisms not only for maintaining mitotic autophagy, but also for the potential chemotherapy resistance in some cancer patients bearing ULK1-ATG13 mutations in CDK1 motifs.

## Materials and Methods

### Antibodies and reagents

The autophagy antibody sampler kit (#4445), ULK1 Antibody Sampler Kit (#8359), Autophagy Induction (ULK1 Complex) Antibody Sampler Kit (#46486), the cell cycle regulation antibody sampler kit II (#9870), Phospho-(Ser) Kinase Substrate Antibody Sampler Kit (#9615), Phospho-Threonine-Proline Mouse mAb (P-Thr-Pro-101) (#9391), anti-ULK1 (#4776) antibody, the HRP-linked anti-rabbit and anti-mouse IgG antibodies were all from Cell signaling technology. The anti-FLAG (F3165) antibody was acquired from Sigma and anti-β-Tubulin, anti-GAPDH and anti-β-Actin antibodies from Beijing TransGen Biotech (Beijing, China). The GlutaMAX supplement and puromycin dihydrochloride were from Gibco. The secondary fluorescently conjugated antibodies, anti-fade prolong Gold with DAPI were from Molecular Probes. Prestained Protein Ladder (26616) and M-PER buffer were from Thermo Pierce. NH_4_Cl, RO-3306 and Thymidine were from Sigma. Nocodazole and SBI-0206965 were from Selleckchem. FITC Annexin V Apoptosis Detection Kit I (#556547) was from BD Biosciences. Methyl cellulose (#69016260) was from Sinopharm Chemical Reagent Co., Ltd. Matrigel Matrix (#354234) was from BD. Protease inhibitor and phosphatase inhibitor cocktails were from Roche and the PVDF membrane from Millipore.

### Cell culture and stable cell lines establishment

HeLa, HCT 116, RPE1 and HEK-293T cells were all cultured in DMEM medium (without L-Glutamine) supplemented with 10% FBS, 2 mM GlutaMAX and 1% penicillin/streptomycin (P/S). The plasmid for pBobi-FLAG-mULK1 contains one FLAG tag and the affinity for FLAG antibody was lower than ULK1 antibody. Therefore, in order to enhance its affinity to FLAG antibody, 3×FLAG was added to mULK1 C-terminus [67]. Stable cell lines were constructed as described previously [68]. HEK-293T or HeLa cells stably expressing mULK1-3×FLAG were maintained in DMEM complete medium containing 1 μg/ml puromycin.

### CRISPR/Cas9 technology

The gRNA targeted to human ULK1 was designed with CRISPR Design (http://crispr.mit.edu). The sequence (human ULK1: 5’-GCCCTTGAAGACCACCGCGA-3’; human ATG13: 5’-CACATGGACCTCCCGACTGC-3’) was selected and subcloned into PX458 vector. HeLa and HEK-293T cells were transiently transfected with PX458-ULK1/ATG13-gRNA with Fugene 6. The cells were diluted into 0.5 cell/100 μL at 96-well plate after 12 h transfection. Single cell in 96-well-plate was cultured in DMEM complete medium to form single-cell-clone that was cultured in 24-well-plate and subjected to immunoblotting analysis using ULK1 or ATG13 specific antibody. One of single-cell-clone could not be detected with ULK1/ATG13 antibody was selected as ULK1/ATG13-knockout cell. ULK1 and ATG13 double knockout cells were established based on ULK1-knockout cells using PX458-ATG13-gRNA.

### Immunoprecipitation and Western blots

The procedure was instructed as previously [68]. Most immunoprecipitation experiments were conducted in HEK-293T derived cell lines due to higher expression level for the exogenous protein. Briefly, HEK-293T cells stably expressing GFP-3FLAG or mULK1-3FLAG were lysed with M-PER supplemented with protease inhibitors and phosphatase inhibitors and centrifuged at 4°C 14,000g for 10 min. The supernatant mixed with pre-incubated Protein G Dynabeads and FLAG antibody at 4°C for 12 h and washed three times with lysis buffer. Then the immunoprecipitate was denatured in 1×SDS-PAGE buffer at 95°C for 7 min and subjected to SDS-PAGE and immunoblotting or Coomassie brilliant blue staining.

### Apoptosis by flow cytometry

Cells treated with Taxol combine with or without chloroquine were subjected to apoptosis assay according to the manufacturer instructions (BD Biosciences). Briefly, cells were trypsinized and washed twice with ice cold PBS, and stained with PI and/or FITC-Annexin V in apoptosis buffer for 30 min. The FITC and PE channel were used for flow cytometry detection and the data were analyzed by FlowJo.

### Cell cycle synchronization

Various cell synchronization methods are used in this paper, which have been used previously. Briefly, a double-thymidine (2.5 μM) block arrested cells in G1/S border and cells progress through S, G2 and M phase after release. A double-thymidine or single-thymidine block in combination with nocodazole (100 ng/mL) or STLC (5 μM) treatment arrested cells in prometaphase or prophase. A double-thymidine block in combination with RO-3306 (10 μM) treatment arrested cells in late G2 phase and progressed into mitosis after RO-3306 washout for three times with prewarmed PBS.

### Cell cycle analysis by flow cytometry/FACS

Cells for cell cycle analysis were trypsinized with 0.25% Trypsin/EDTA. After washing with ice-cold PBS twice, cells were fixed with −20°C 75% ethanol overnight and then stained with PI/RNase staining buffer (BD pharmingen) for 15 minutes at room temperature and analyzed with flow cytometry (Beckman Coulter, Cytoflex). Alternatively, for mitotic index analysis, cells fixed were stained with phospho-Histone H3 (S10) at 1:1600 for 2 hours at room temperature and washed twice before Alexa-488 conjugated anti-rabbit IgG staining. After washing twice, PI/RNase staining was conducted as described above before flow cytometry analysis. The data were analyzed by ModFit LT 4.1 and Flow Jo 7.6 software.

### Lambda Phosphatase treatment

IP products from mULK1-3FLAG overexpressing HEK293T cells were aliquoted and treated with reaction buffer, reaction buffer containing 1 μL (400 units) lambda phosphatase (P0753S, NEB), reaction buffer containing 1 μL lambda phosphatase plus 1×phosphatase inhibitors cocktail (Roche) in at 30°C for 30 min with gently shaking. Then the reaction products were denatured at 95°C for 7 min and subjected to immunoblotting with FLAG or Serine/Threonine specific antibody.

### Mass spectrometry

For mass spectrometry assays, immunoprecipitates using FLAG antibody from asynchronous or mitotic 293T cells expressing FLAG-tagged mULK1/ATG13 were separated by SDS-PAGE, the gel was stained with Coomassie brilliant blue, and the FLAG-tagged mULK1/ATG13 band in each lane was excised. Samples were subjected to mass spectrometry analysis for mULK1/ATG13 phosphorylation by Core Facility Center for Life Sciences, University of Science and Technology of China, School of Life Science and Technology, ShanghaiTech University. The identified phosphorylation sites specific in mitotic cells were selected as the candidates for ULK1/ATG13 phosphorylation sites in mitosis.

### *In vitro* kinase assay

Anti-FLAG immunoprecipitates from asynchronous 293T cells overexpressing FLAG-tagged mULK1 (wild-type or K46I kinase dead) or ATG13 were washed three times with M-PER and then resuspended in ice-cold kinase buffer (50 mM Tris-HCl pH 7.5 @ 25°C, 10 mM MgCl_2_, 0.1 mM EDTA, 2 mM DTT, 0.01% Brij 35). The immunoprecipitates were then incubated with or without 270 ng purified CDK1/cyclin B (Life technologies, Part Number: PV3292, Lot Number: 1816161K) in 20 μL reaction mix (kinase buffer and 20 μM ATP) pretreated with or without 10 μM CDK1 inhibitor RO-3306 at 30°C with constant shaking for 30 min. The reaction was quenched by mixing with 5 μL 5×SDS-sample buffer and boiling at 95°C for 7 min.

### Mouse Model

Four-week-old female BALB/c nude mice were purchased from Nanjing Biomedical Research Institute of Nanjing University (Nanjing, China). All mice were kept in an animal room under the specific-pathogen-free (SPF) condition. The mice were fed with sterilized food and autoclaved tap water freely. The protocol involving animals was approved by the ethical and humane committee of Hefei Institutes of Physical Science, Chinese Academy of Sciences and carried out strictly in accordance with the related regulations (Hefei, China). After one week, HeLa/HeLa-ATG13 knockout/HeLa-ULK1&ATG13 double knockout cells (1 × 10^7^) suspended in 100 μL PBS/Matrigel Matrix (1:1) were injected into the subcutaneous space on the right flank of BALB/c nude mice. The mice bearing HeLa/HeLa-ATG13 knockout cells were randomly divided into two groups of five mice each seven days post implantation. Mice were intraperitoneally injected with 0.5% (M/V) methyl cellulose used as the vehicle solution or SBI-0206965 in 0.5% methyl cellulose (20 mg/kg/d) every day for 33 days. Tumor growth was evaluated every day and tumor volume was calculated as: volume = 1/2(length × width^2^). At the end of the experiment, the mice were sacrificed by cervical dislocation and tumors were harvested and weighted up.

### Quantification and Statistical Analysis

ImageJ software was used to quantify the relative protein value for Western Blot band and Graphpad prism 6 was used to analyze the data using Student’s *t*-test for two groups. P values < 0.05 were considered as statistically significant.

## Acknowledgments

We thank Professor Wenchao Wang at High magnetic field laboratory, Chinese Academy of Sciences for providing technical guidance for CRISPR/Cas9, Professor Shengcai Lin at Xiamen University for providing the plasmid of FLAG-mULK1, Professor Zhenye Yang at University of Science and Technology of China for providing the inhibitors of Hesperadin, MLN8237 and STLC, Professor Qingsong Liu at High magnetic field laboratory for providing the inhibitors of GSK1070916, AZD1152-HQPA, PHA793887 and AZD5438, Dr. Piliang Hao at ShanghaiTech University and Dr. Gao Wu at University of Science and Technology of China for mass spectrometry analysis. This work was supported by the National Key Research and Development Program of China (#2016YFA0400900), Major/Innovative Program of Development Foundation of Hefei Center for Physical Science and Technology (2016FXCX004), Key Program of 13^th^ five-year plan, CASHIPS (KP-2017-26) and CASHIPS Director’s Fund (YZJJ201704) to Xin Zhang, CASHIPS Director’s Fund (YZJJ2019QN17) and Anhui Natural Science Foundation (1908085MC88) to Zhiyuan Li.

## Author Contributions

Z.L. performed most of the experiments; X.T. performed the mouse model assays; X.J. performed the apoptosis detection by flow cytometry; D.W. counted the cell number of HeLa/HeLa-DKO; Z.L. and X.Z. designed the experiments, analyzed data and wrote the paper. All authors edited the manuscript.

## Supporting information

**Fig. S1.**
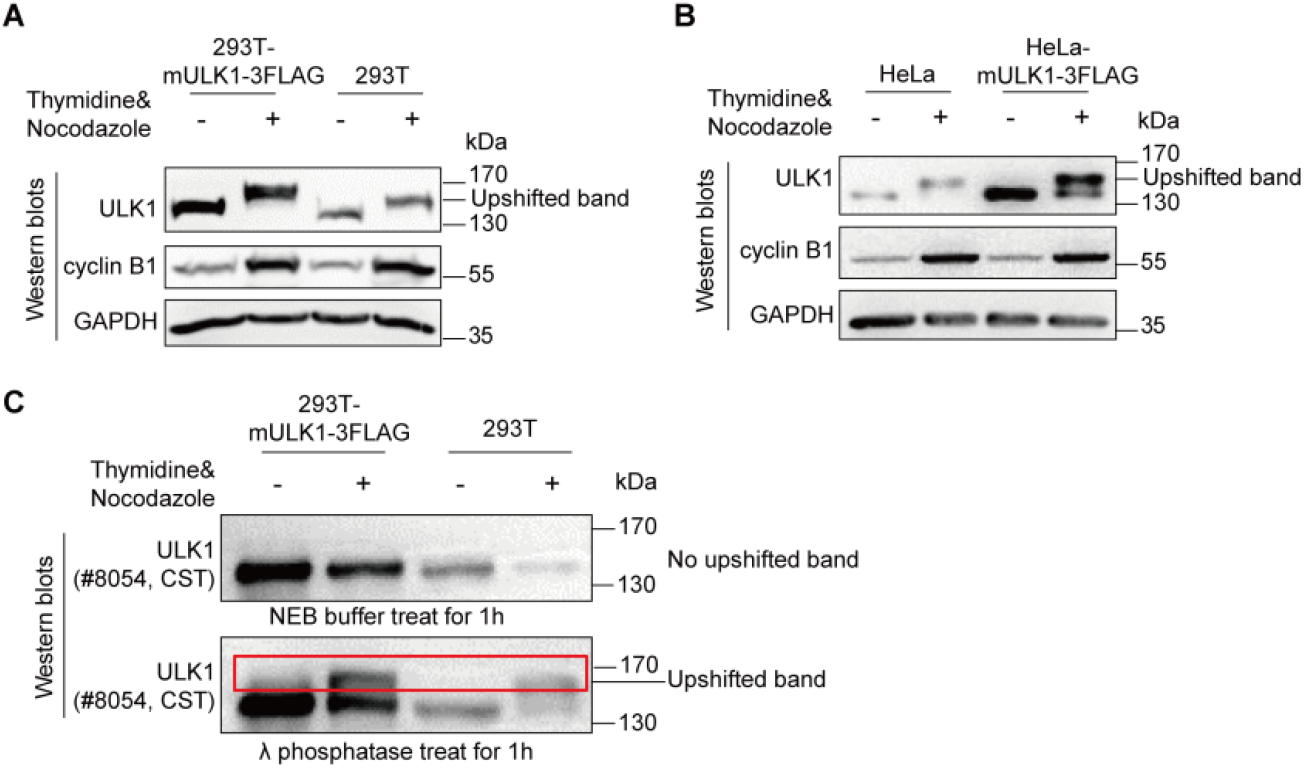
ULK1 is phosphorylated and upshifted in mitosis. (A-B) Both endogenous human and exogenous mouse ULK1 are upshifted in thymidine and nocodazole-arrested mitosis. 293T and HeLa cells with or without FLAG-tagged mULK1 overexpression were synchronized into mitosis by single-thymidine and nocodazole for Western blots analysis. (C) ULK1 phosphorylation in mitosis interferes with a ULK1 antibody recognition. The ULK1 antibody (Cell signaling technology, #8054) could not recognize the upshifted band for mitotic ULK1 but could recognize when the PVDF membrane was treated with lambda phosphatase for 1 h.

**Fig. S2.**
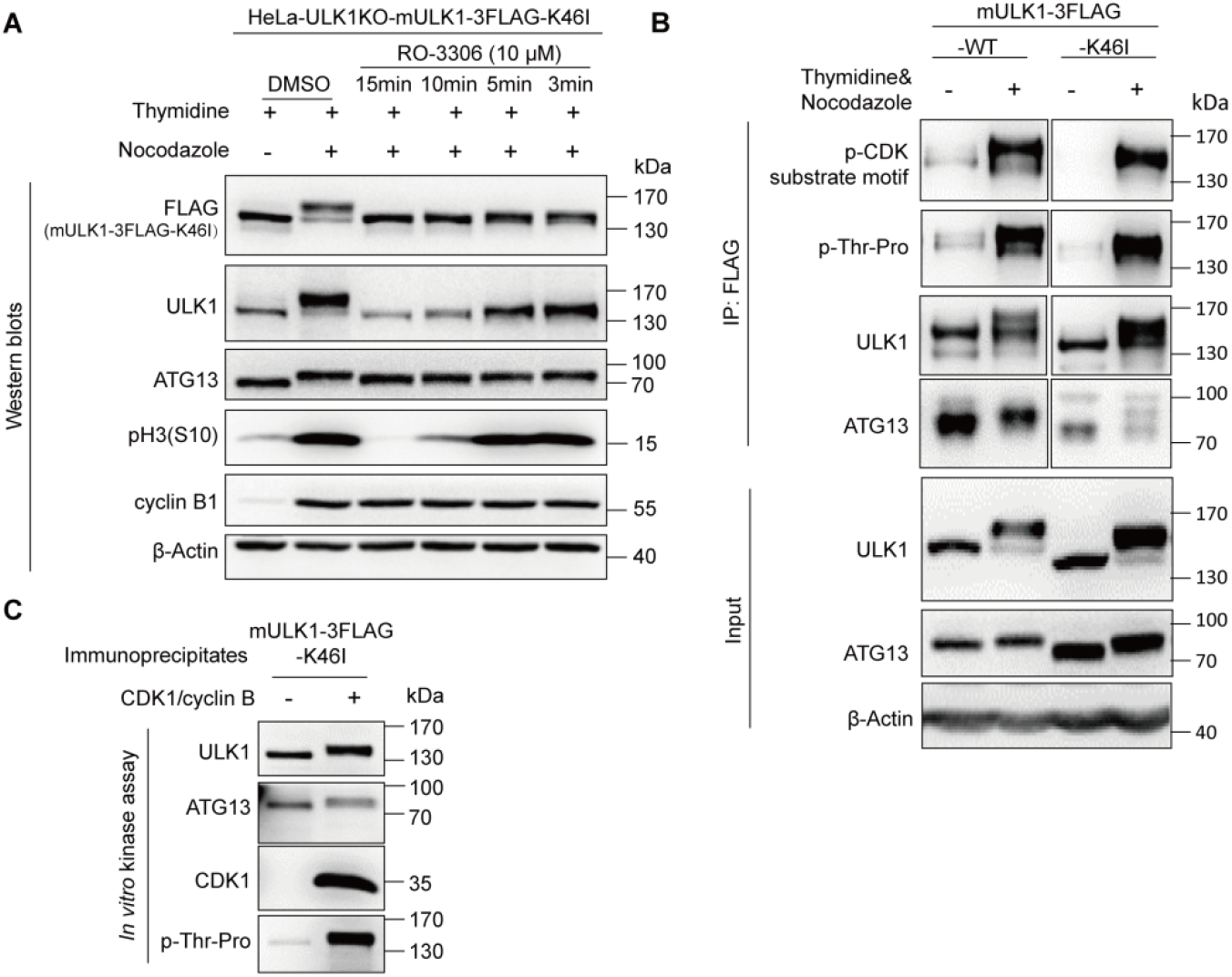
CDK1 regulates ULK1 phosphorylation independent of ULK1 kinase activity. (A-B) K46I kinase dead ULK1 also underwent significant electrophoretic mobility shift and phosphorylation in mitosis as wild-type ULK1. HeLa-ULK1 knockout cells reconstituted with FLAG-tagged mULK1-K46I or -wild type were treated as Figure 2A and 3B (the lower panel) and then analyzed by Western blots and immunoprecipitation respectively. (C) In vitro kinase assay indicated that purified CDK1/cyclin B could induce K46I kinase dead ULK1 undergo significant electrophoretic mobility shift and phosphorylation.

**Fig. S3.**
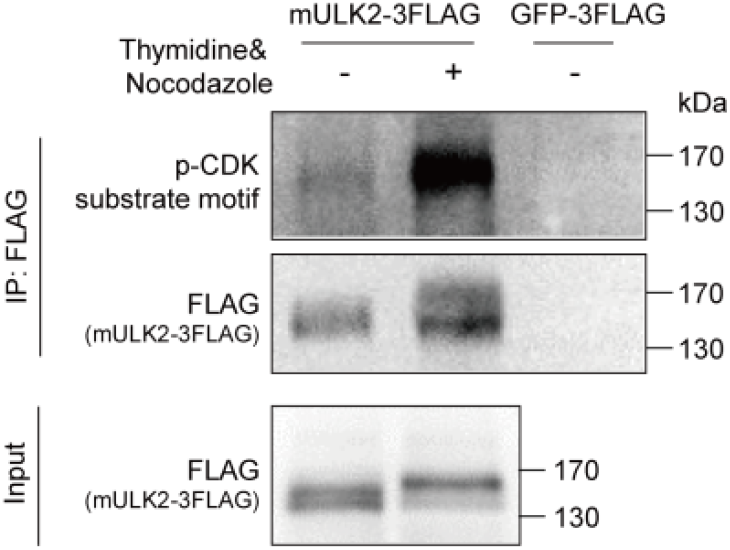
ULK2 was also phosphorylated and upshifted in mitosis. 293T cells transiently transfected with FLAG-tagged mULK2 treated as Figure 3A were analyzed by Western blots and immunoprecipitation.

**Fig. S4.**
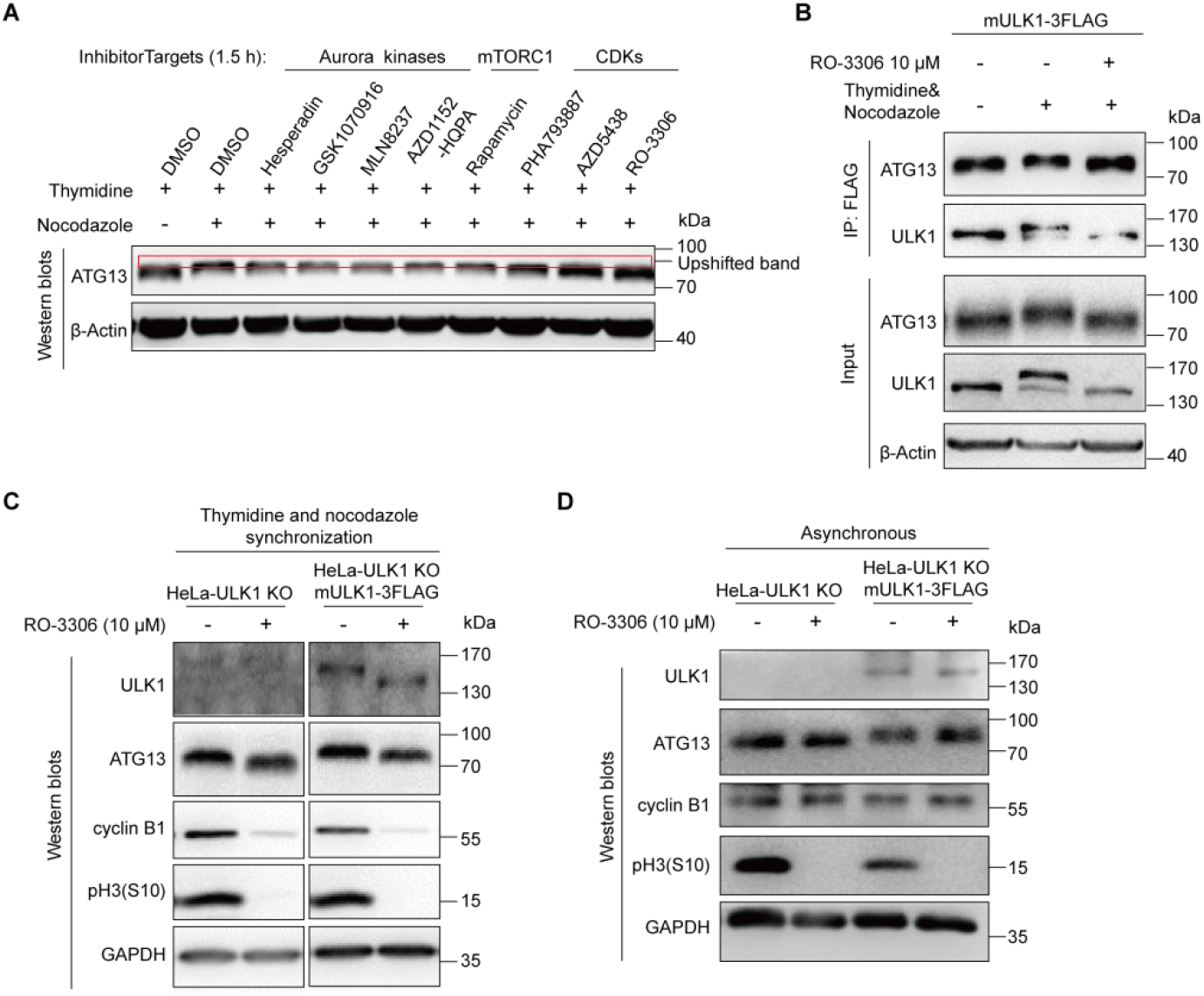
ATG13 is upshifted in mitosis. (A) HeLa cells treated as Figure 4B (the upper panel) for ATG13 mobility shift analysis. (B) ATG13 mobility shift in mitosis is decreased by CDK1 inhibitor RO-3306, similarly to ULK1, although to a less extent. 293T cells overexpressing FLAG-tagged mULK1 were synchronized by single-thymidine in the presence or absence of nocodazole, treated with 10 μM RO-3306 for 5 or 30 min. The co-immunoprecipitate by FLAG antibody was subjected to immunoblotting with ATG13, FIP200 antibodies. (C-D) ULK1 expression level does not affect ATG13 mobility shift in mitosis. HeLa ULK1-knockout cells with or without FLAG-tagged mULK1 expression synchronized into mitosis with thymidine and nocodazole (C) or in asynchrounous condition (D) were treated with 10 μM RO-3306 for 30 min for Western blots analysis.

**Fig S5.**
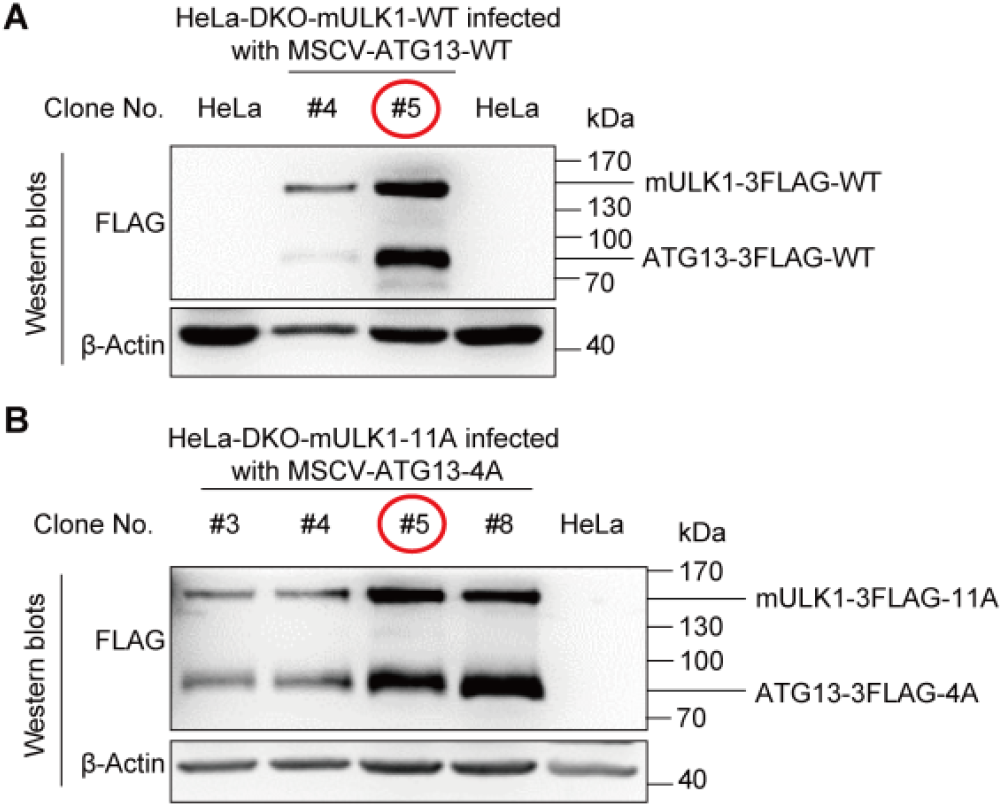
Establishment of ULK1 and ATG13 double wild-type or mutant cell line. The cell lines were established based on HeLa-DKO cell reconstituted with FLAG-tagged wild-type or 11A mutant mULK1 and the method is the same as HeLa-ULK1/ATG13 knockout cell line establishment shown in Figure S7A and S9A.

**Fig. S6.**
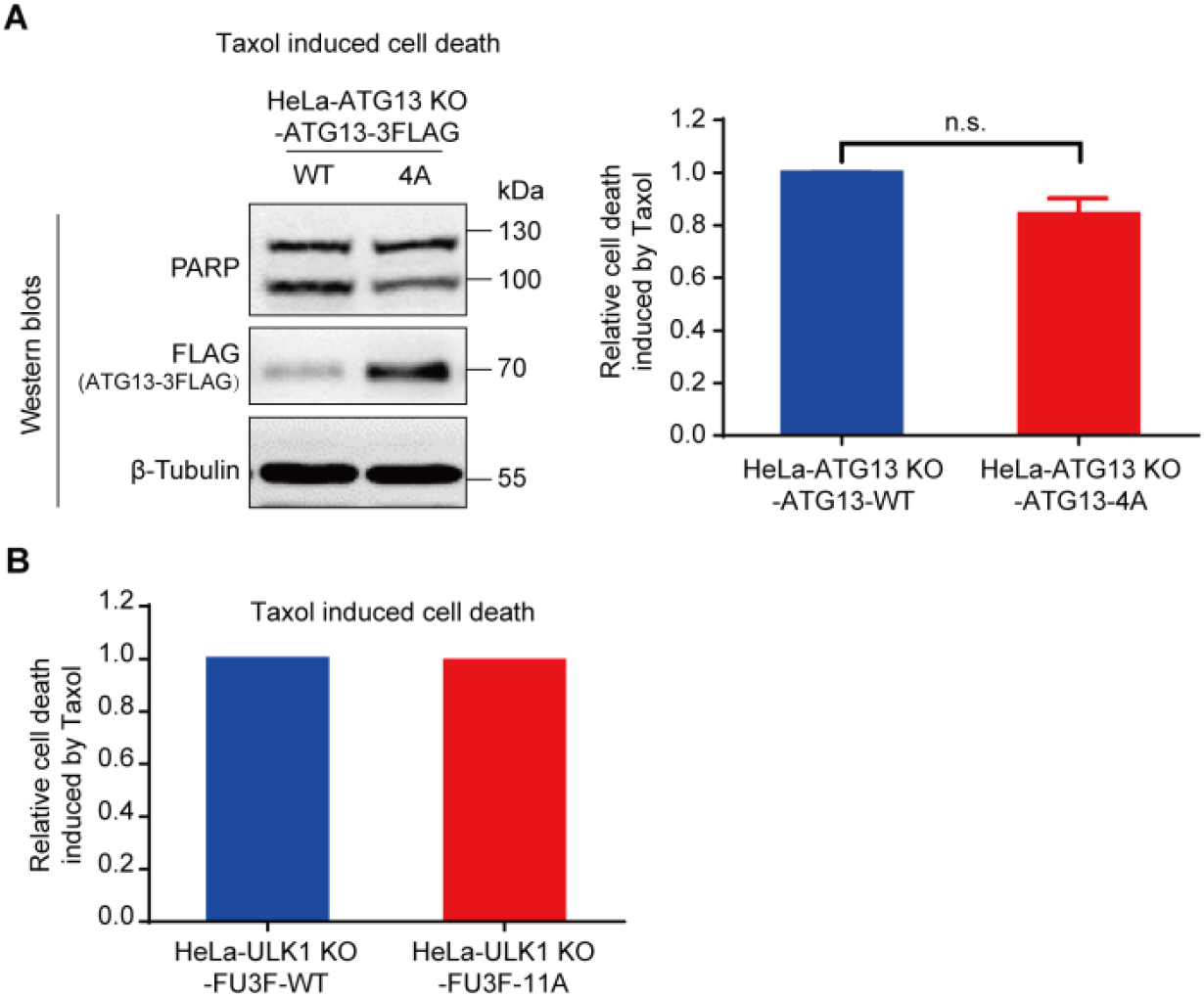
Either ATG13 or ULK1 mutant has mild effect on Taxol-induced cell death. HeLa-ULK1 knockout cells reconstituted with FLAG-tagged wild-type or 11A mutant mULK1 or HeLa-ATG13 knockout cells reconstituted with FLAG-tagged wild-type or 4A mutant ATG13 were treated as Figure 6B were analyzed by Western blots and/or flow cytometry. n=3, n.s., not significant.

**Fig S7.**
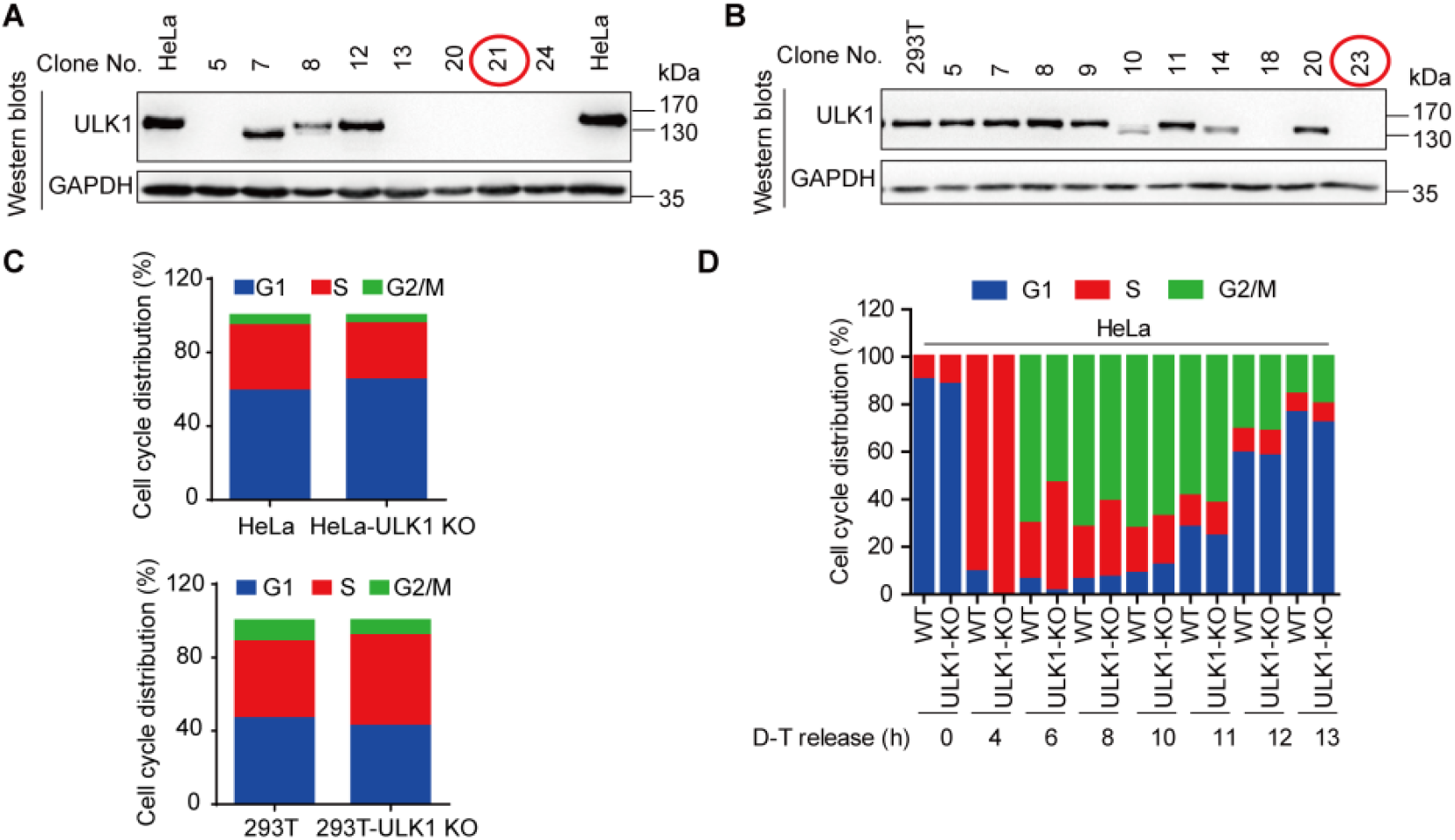
ULK1-knockout cells were established and ULK1 knockout slightly delays S/G2 transition. (A-B) ULK1-knockout cells establishment. HeLa and 293T cells transiently transfected with the CRISPR/Cas9 plasmid subcloned gRNA for human ULK1 were screened by Western blots and the ULK1-knockout clones were identified. The red circles indicate the ULK1-knockout clones for the following assay. (C) ULK1 knockout does not affect cell cycle distribution in HeLa and 293T cells. Cell cycle distribution was analyzed by flow cytometry in asynchronous wild-type and ULK1-knockout HeLa or 293T cells. (D) ULK1 knockout slightly delays S/G2 transition. HeLa wild-type or ULK1-knockout cells synchronized with double-thymidine and nocodazole were subjected to cell cycle analysis by flow cytometry.

**Fig. S8.**
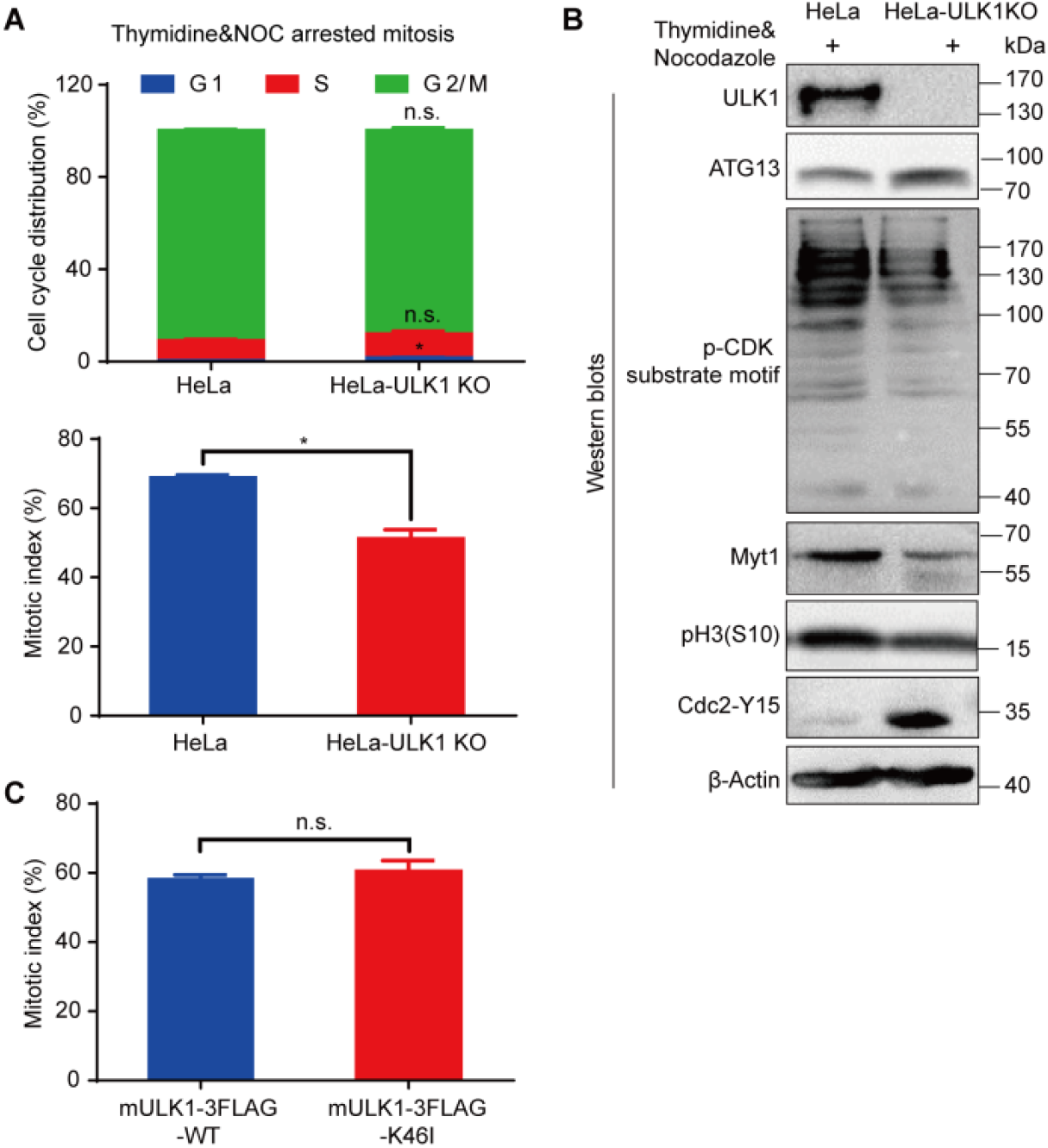
Mitotic progression was delayed in ULK1 knockout but not K46I-kinase dead ULK1 cell line. (A-B) Mitotic index is decreased in ULK1-knockout cells synchronized by single-thymidine and nocodazole. HeLa wild-type or ULK1-knockout cells synchronized into mitosis were subjected to either PI and pH3(S10) co-staining for cell cycle and mitotic index analysis by flow cytometry (A) or Western blots analysis for cell cycle markers (B). (C) Mitotic progression was not affected by K46I-kinase dead ULK1. HeLa ULK1-knockout cells reconstituted with FLAG-tagged wild-type or K46I kinase dead mULK1 were synchronized into mitosis and subjected to pH3(S10) staining for mitotic index analysis by flow cytometry. n=3, n.s., not significant, *p < 0.05.

**Fig. S9.**
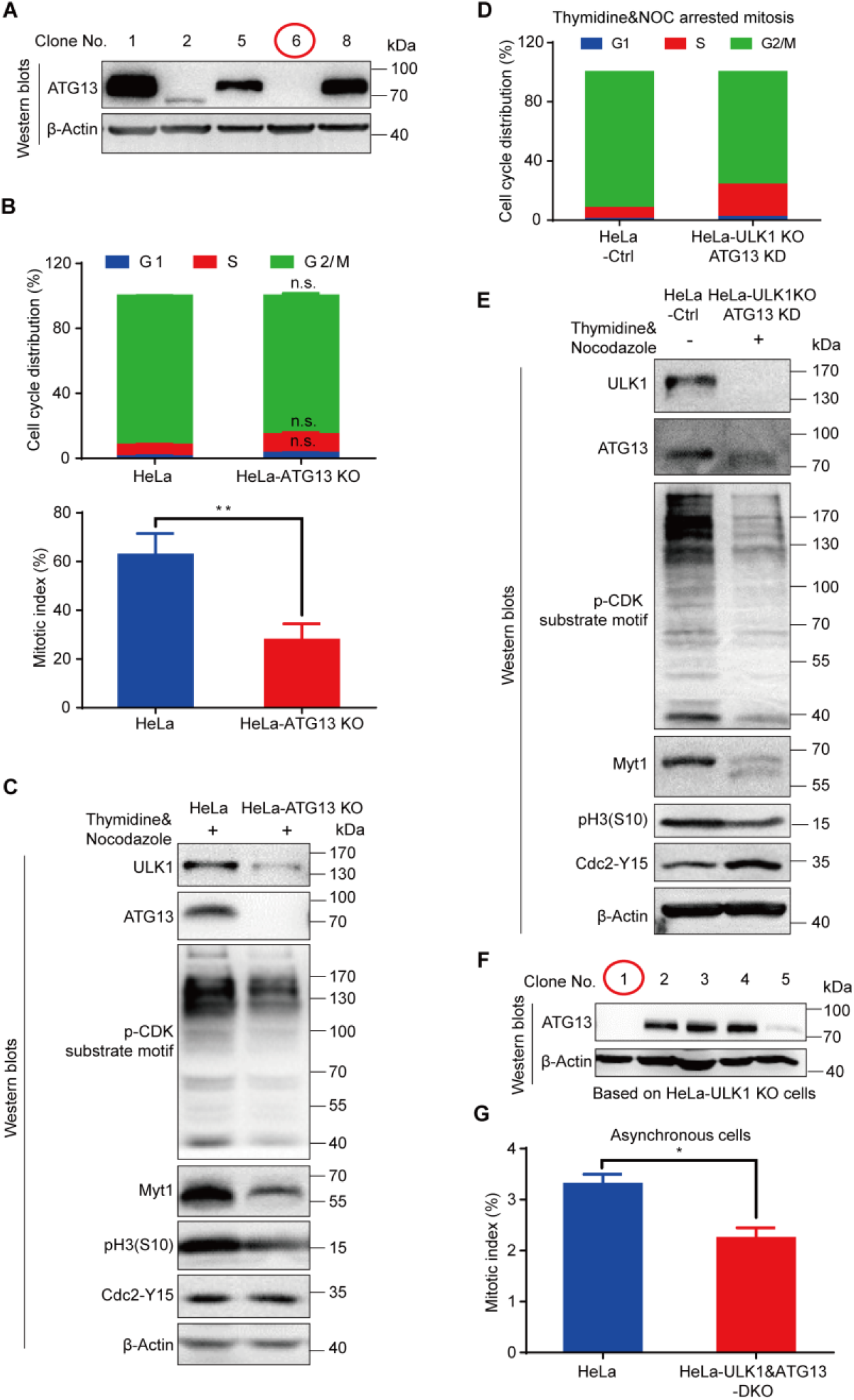
ULK1 and ATG13 coordinate to regulate cell cycle progression. (A, F) ATG13-knockout or ULK1 and ATG13 double knockout cells establishment. HeLa wild-type or ULK1-knockout cells transiently transfected with the CRISPR/Cas9 plasmid subcloned gRNA for human ATG13 were screened by Western blots and the ATG13-knockout clones were identified. The red circles indicate the ATG13-knockout or ULK1 and ATG13 double knockout clones for the following assay. (B-C) Mitotic index was decreased in ATG13-knockout cells synchronized by single-thymidine and nocodazole. HeLa wild-type or ATG13-knockout cells synchronized into mitosis were subjected to PI and pH3(S10) co-staining for cell cycle and mitotic index analysis by flow cytometry (B) or Western blots analysis for cell cycle markers (C). (D) ULK1 knockout combined with ATG13 “knockdown” inhibits S/G2 transition. HeLa ULK1-knockout cells transiently transfected with the CRISPR/Cas9 vector control or plasmid subcloned gRNA for human ATG13 were synchronized with thymidine and nocodazole for cell cycle analysis by flow cytometry. (E) The cell lysate collected from (D) was subjected to Western blots by indicated antibodies. (G) Double knockout ULK1 and ATG13 decreases mitotic index. HeLa wild-type or ULK1&ATG13-double knockout cells were subjected to pH3(S10) staining for mitotic index analysis by flow cytometry. n=3, n.s., not significant, *p < 0.05, **p < 0.01.

**Fig. S10.**
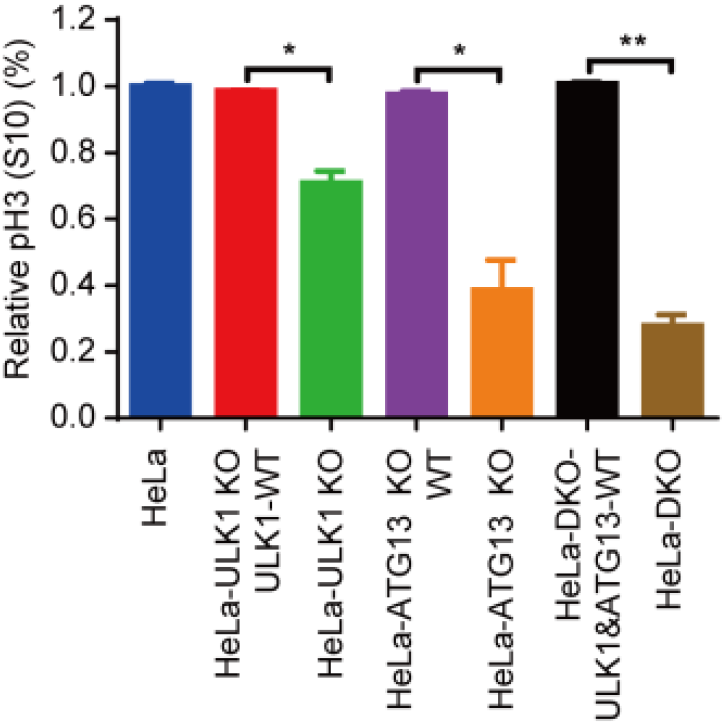
Relative mitotic index in knockout cells reconstituted with or without FLAG-tagged mULK1 and/or ATG13. Cells synchronized by thymidine and nocodazole were subjected to PI and pH3(S10) co-staining for cell cycle and mitotic index analysis by flow cytometry. n=3, *p < 0.05, **p < 0.01.

**Fig. S11.**
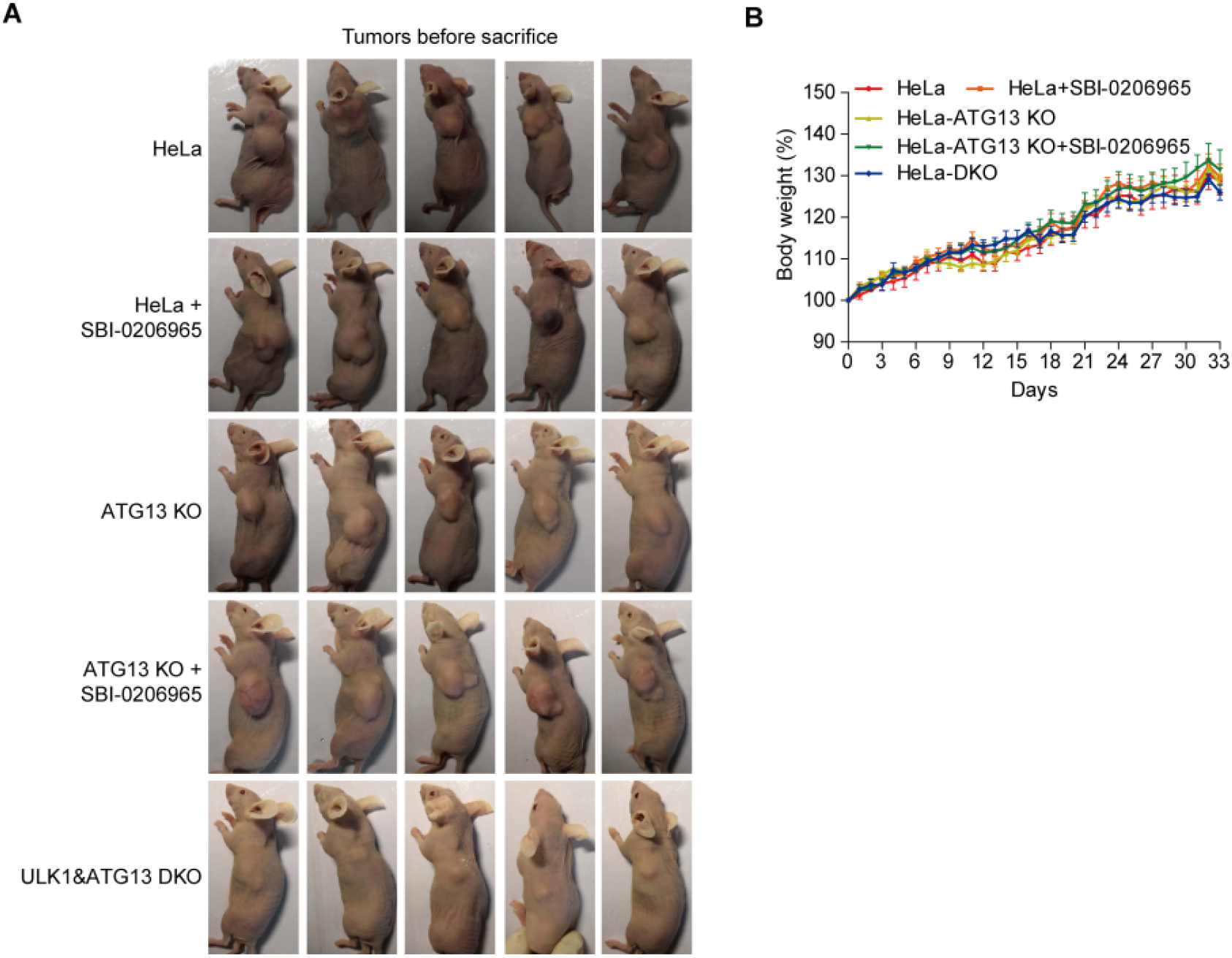
Mouse model. (A) The mouse model in nude mice bearing wild-type or knockout cells treated with or without ULK1 kinase inhibitor SBI-0206965 was established. (B) Time course of the body weight for nude mice in (A).

## References

1. Suzuki, H., et al., Structural biology of the core autophagy machinery. Current Opinion in Structural Biology, 2017. 43: p. 10–17.

2. Levine, B. and G. Kroemer, Biological Functions of Autophagy Genes: A Disease Perspective. Cell, 2019. 176(1-2): p. 11–42.

3. Klionsky, D.J., et al., Guidelines for the use and interpretation of assays for monitoring autophagy. Autophagy, 2016. 12(1): p. 1–222.

4. Lee, I.H., et al., Atg7 modulates p53 activity to regulate cell cycle and survival during metabolic stress. Science, 2012. 336(6078): p. 225–228.

5. Melkoumian, Z.K., et al., Mechanism of cell cycle regulation by FIP200 in human breast cancer cells. Cancer Res, 2005. 65(15): p. 6676–84.

6. Fré mont, S., et al., Beclin - 1 is required for chromosome congression and proper outer kinetochore assembly. EMBO reports, 2013. 14(4): p. 364–372.

7. Maskey, D., et al., ATG5 is induced by DNA-damaging agents and promotes mitotic catastrophe independent of autophagy. Nature communications, 2013. 4.

8. Egan, D.F., et al., Phosphorylation of ULK1 (hATG1) by AMP-activated protein kinase connects energy sensing to mitophagy. Science, 2011. 331(6016): p. 456–461.

9. Kim, J., et al., AMPK and mTOR regulate autophagy through direct phosphorylation of Ulk1. Nature Cell Biology, 2011. 13(2): p. 132–U71.

10. Egan, D.F., et al., Small Molecule Inhibition of the Autophagy Kinase ULK1 and Identification of ULK1 Substrates. Mol Cell, 2015. 59(2): p. 285–97.

11. Jeong, Y.-T., et al., The ULK1-FBXW5-SEC23B nexus controls autophagy. eLife, 2018. 7: p. e42253.

12. Li, Z. and X. Zhang, Kinases Involved in Both Autophagy and Mitosis. International Journal of Molecular Sciences, 2017. 18(9): p. 1884.

13. Li, Z. and X. Zhang, Autophagy in mitotic animal cells. Science Bulletin, 2015: p. 1–3.

14. Mathiassen, S.G., D. De Zio, and F. Cecconi, Autophagy and the Cell Cycle: A Complex Landscape. Front Oncol, 2017. 7: p. 51.

15. Li, Z., et al., Autophagic flux is highly active in early mitosis and differentially regulated throughout the cell cycle. Oncotarget, 2016. 7(26): p. 39705.

16. Furuya, T., et al., Negative regulation of Vps34 by Cdk mediated phosphorylation. Mol Cell, 2010. 38(4): p. 500–11.

17. Liu, L., et al., Robust autophagy/mitophagy persists during mitosis. Cell Cycle, 2009. 8(10): p. 1616–1620.

18. Doménech, E., et al., AMPK and PFKFB3 mediate glycolysis and survival in response to mitophagy during mitotic arrest. Nature cell biology, 2015. 17(10): p. 1304.

19. Puente, C., R.C. Hendrickson, and X. Jiang, Nutrient-regulated phosphorylation of ATG13 inhibits starvation-induced autophagy. Journal of Biological Chemistry, 2016. 291(11): p. 6026–6035.

20. Shang, L., et al., Nutrient starvation elicits an acute autophagic response mediated by Ulk1 dephosphorylation and its subsequent dissociation from AMPK. Proceedings of the National Academy of Sciences, 2011. 108(12): p. 4788–4793.

21. Vassilev, L.T., et al., Selective small-molecule inhibitor reveals critical mitotic functions of human CDK1. Proceedings of the National Academy of Sciences, 2006. 103(28): p. 10660–10665.

22. Gwinn, D.M., J.M. Asara, and R.J. Shaw, Raptor is phosphorylated by cdc2 during mitosis. PloS one, 2010. 5(2): p. e9197.

23. Charrasse, S., et al., Ensa controls S-phase length by modulating treslin levels. Nature communications, 2017. 8(1): p. 206.

24. Contreras-Vallejos, E., et al., Searching for novel Cdk5 substrates in brain by comparative phosphoproteomics of wild type and Cdk5−/− mice. PloS one, 2014. 9(3): p. e90363.

25. Zhang, H., et al., An Eya1-Notch axis specifies bipotential epibranchial differentiation in mammalian craniofacial morphogenesis. eLife, 2017. 6.

26. Ganley, I.G., et al., ULK1· ATG13· FIP200 complex mediates mTOR signaling and is essential for autophagy. Journal of Biological Chemistry, 2009. 284(18): p. 12297–12305.

27. Jung, C.H., et al., ULK-Atg13-FIP200 complexes mediate mTOR signaling to the autophagy machinery. Molecular biology of the cell, 2009. 20(7): p. 1992–2003.

28. Lin, S.Y., et al., GSK3-TIP60-ULK1 signaling pathway links growth factor deprivation to autophagy. Science, 2012. 336(6080): p. 477–81.

29. Hosokawa, N., et al., Nutrient-dependent mTORC1 association with the ULK1-Atg13-FIP200 complex required for autophagy. Mol Biol Cell, 2009. 20(7): p. 1981–91.

30. Skoufias, D.A., et al., S-trityl-L-cysteine is a reversible, tight binding inhibitor of the human kinesin Eg5 that specifically blocks mitotic progression. J Biol Chem, 2006. 281(26): p. 17559–69.

31. Adams, N.D., et al., Discovery of GSK1070916, a potent and selective inhibitor of Aurora B/C kinase. J Med Chem, 2010. 53(10): p. 3973–4001.

32. Walsby, E., et al., Effects of the aurora kinase inhibitors AZD1152-HQPA and ZM447439 on growth arrest and polyploidy in acute myeloid leukemia cell lines and primary blasts. Haematologica, 2008. 93(5): p. 662–9.

33. Gorgun, G., et al., A novel Aurora-A kinase inhibitor MLN8237 induces cytotoxicity and cell-cycle arrest in multiple myeloma. Blood, 2010. 115(25): p. 5202–13.

34. Hauf, S., et al., The small molecule Hesperadin reveals a role for Aurora B in correcting kinetochore–microtubule attachment and in maintaining the spindle assembly checkpoint. The Journal of Cell Biology, 2003. 161(2): p. 281–294.

35. Thoreen, C.C. and D.M. Sabatini, Rapamycin inhibits mTORC1, but not completely. Autophagy, 2009. 5(5): p. 725–6.

36. Byth, K.F., et al., AZD5438, a potent oral inhibitor of cyclin-dependent kinases 1, 2, and 9, leads to pharmacodynamic changes and potent antitumor effects in human tumor xenografts. Mol Cancer Ther, 2009. 8(7): p. 1856–66.

37. Brasca, M.G., et al., Optimization of 6,6-dimethyl pyrrolo[3,4-c]pyrazoles: Identification of PHA-793887, a potent CDK inhibitor suitable for intravenous dosing. Bioorg Med Chem, 2010. 18(5): p. 1844–53.

38. Bayliss, R., et al., On the molecular mechanisms of mitotic kinase activation. Open biology, 2012. 2(11): p. 120136–120136.

39. Chan, E.Y.W., et al., Kinase-inactivated ULK proteins inhibit autophagy via their conserved C-terminal domains using an Atg13-independent mechanism. Molecular and cellular biology, 2009. 29(1): p. 157–171.

40. Shain, A.H., et al., Exome sequencing of desmoplastic melanoma identifies recurrent NFKBIE promoter mutations and diverse activating mutations in the MAPK pathway. Nat Genet, 2015. 47(10): p. 1194–9.

41. Hoadley, K.A., et al., Cell-of-Origin Patterns Dominate the Molecular Classification of 10,000 Tumors from 33 Types of Cancer. Cell, 2018. 173(2): p. 291–304.e6.

42. Robertson, A.G., et al., Comprehensive Molecular Characterization of Muscle-Invasive Bladder Cancer. Cell, 2017. 171(3): p. 540–556.e25.

43. Hugo, W., et al., Genomic and Transcriptomic Features of Response to Anti-PD-1 Therapy in Metastatic Melanoma. Cell, 2016. 165(1): p. 35–44.

44. The Cancer Genome Atlas Research, N., et al., Comprehensive molecular profiling of lung adenocarcinoma. Nature, 2014. 511: p. 543.

45. Campbell, J.D., et al., Distinct patterns of somatic genome alterations in lung adenocarcinomas and squamous cell carcinomas. Nat Genet, 2016. 48(6): p. 607–16.

46. Giannakis, M., et al., Genomic Correlates of Immune-Cell Infiltrates in Colorectal Carcinoma. Cell Rep, 2016. 15(4): p. 857–865.

47. The Cancer Genome Atlas Research, N., et al., Comprehensive molecular characterization of clear cell renal cell carcinoma. Nature, 2013. 499: p. 43.

48. Sato, Y., et al., Integrated molecular analysis of clear-cell renal cell carcinoma. Nat Genet, 2013. 45(8): p. 860–7.

49. Ciriello, G., et al., Comprehensive Molecular Portraits of Invasive Lobular Breast Cancer. Cell, 2015. 163(2): p. 506–19.

50. Levy, J.M., et al., Autophagy inhibition improves chemosensitivity in BRAF(V600E) brain tumors. Cancer Discov, 2014. 4(7): p. 773–80.

51. Gao, M., Y. Xu, and L. Qiu, Enhanced combination therapy effect on paclitaxel-resistant carcinoma by chloroquine co-delivery via liposomes. Int J Nanomedicine, 2015. 10: p. 6615–32.

52. Levy, J.M., et al., Autophagy inhibition overcomes multiple mechanisms of resistance to BRAF inhibition in brain tumors. Elife, 2017. 6.

53. Brito, D.A., Z. Yang, and C.L. Rieder, Microtubules do not promote mitotic slippage when the spindle assembly checkpoint cannot be satisfied. The Journal of Cell Biology, 2008. 182(4): p. 623–629.

54. Hirota, T., et al., Histone H3 serine 10 phosphorylation by Aurora B causes HP1 dissociation from heterochromatin. Nature, 2005. 438: p. 1176.

55. Shuda, M., et al., CDK1 substitutes for mTOR kinase to activate mitotic cap-dependent protein translation. Proceedings of the National Academy of Sciences, 2015. 112(19): p. 5875–5882.

56. Potapova, T.A., et al., Fine Tuning the Cell Cycle: Activation of the Cdk1 Inhibitory Phosphorylation Pathway during Mitotic Exit. Molecular Biology of the Cell, 2009. 20(6): p. 1737–1748.

57. Sorokina, I.V., et al., Involvement of autophagy in the outcome of mitotic catastrophe. Scientific Reports, 2017. 7(1): p. 14571.

58. Fava, L.L., et al., Beclin 1 is dispensable for chromosome congression and proper outer kinetochore assembly. EMBO reports, 2015. 16(10): p. 1233–1236.

59. Joachim, J., et al., Activation of ULK kinase and autophagy by GABARAP trafficking from the centrosome is regulated by WAC and GM130. Molecular cell, 2015. 60(6): p. 899–913.

60. Linares, J.F., et al., Phosphorylation of p62 by cdk1 controls the timely transit of cells through mitosis and tumor cell proliferation. Molecular and cellular biology, 2011. 31(1): p. 105–117.

61. Caballe, A., et al., ULK3 regulates cytokinetic abscission by phosphorylating ESCRT-III proteins. Elife, 2015. 4: p. e06547.

62. Dunlop, E.A., et al., ULK1 inhibits mTORC1 signaling, promotes multisite Raptor phosphorylation and hinders substrate binding. Autophagy, 2011. 7(7): p. 737–47.

63. Randhawa, R., et al., Unc-51 like kinase 1 (ULK1) in silico analysis for biomarker identification: a vital component of autophagy. Gene, 2015. 562(1): p. 40–49.

64. Joo, J.H., et al., Hsp90-Cdc37 chaperone complex regulates Ulk1-and Atg13-mediated mitophagy. Molecular cell, 2011. 43(4): p. 572–585.

65. Burrows, F., H. Zhang, and A. Kamal, Hsp90 activation and cell cycle regulation. Cell Cycle, 2004. 3(12): p. 1530–6.

66. Malumbres, M. and M. Barbacid, Mammalian cyclin-dependent kinases. Trends in biochemical sciences, 2005. 30(11): p. 630–641.

67. Hosokawa, N., et al., Nutrient-dependent mTORC1 association with the ULK1–Atg13–FIP200 complex required for autophagy. Molecular biology of the cell, 2009. 20(7): p. 1981–1991.

68. Li, Z., et al., Ammonia Induces Autophagy through Dopamine Receptor D3 and MTOR. PLoS One, 2016. 11(4): p. e0153526.

